# Beyond profiles: supervised repeat annotation using protein embeddings

**DOI:** 10.64898/2026.05.19.725729

**Authors:** Kaiyu Qiu, Jan Ludwiczak, Andrei N. Lupas, Stanislaw Dunin-Horkawicz

## Abstract

Protein repeats give rise to diverse protein structures and functions, yet their sequence-based detection is often challenging because individual repeat units can diverge substantially in sequence while retaining similar structures. Most sensitive approaches rely on sequence information directly represented by alignments or derived profiles, thereby limiting flexibility and scalability. Here, we introduce TREAD (**<u>T</u>**ransfer learning-based **<u>RE</u>**peat **<u>A</u>**nnotation using Protein Embe**<u>D</u>**dings), a supervised framework that reformulates repeat detection as a residue-level annotation problem and operates directly on embeddings from protein language models. Instead of constructing explicit probabilistic profiles, TREAD learns repeat-specific features implicitly, enabling residue-resolved scoring and flexible repeat segment extraction. Across complementary benchmarks based on RepeatsDB and Pfam, TREAD consistently matches or outperforms the widely used profile-based tool HMMER, particularly in low-data and high-divergence settings. The model exhibits robustness to score thresholding and demonstrates better generalization to independent test sets, including those containing remotely related families. To illustrate the practical utility of TREAD, we applied it to survey β-propeller proteins across the AlphaFold Database and representative proteomes to generate a comprehensive census of this fold. This analysis highlights extensive propeller diversity, identifies lineage-specific expansion patterns across the tree of life, and suggests previously unrecognized relationships between propellers and other repeat folds. Together, TREAD provides a flexible and scalable alternative to profile-based repeat annotation and establishes a general motif-centric framework for protein sequence annotation.

**Author Summary:** Many proteins are assembled from repeated building blocks. These repeats have helped evolution generate a remarkable variety of structures and functions. Yet sensitive detection of individual repeats has long been challenging because their sequences can diverge substantially even when their structures remain similar. Conventional methods often rely on comparing large collections of related sequences to find these patterns. We developed TREAD, a computational method that learns from annotated examples to recognize protein repeats using information captured by protein language models. Rather than building conventional sequence profiles, TREAD identifies repeat units and their boundaries directly from these learned protein representations. In systematic benchmarks, it matched or outperformed HMMER, a widely used profile-based method, particularly when few examples were available or repeat sequences were highly divergent. We then used TREAD to survey β-propellers, proteins assembled from repeated blade-like units, across millions of protein sequences. We uncovered substantial unexplored diversity, differences in their abundance and associated functions across organisms, and unexpected similarities to a previously unclassified barrel-like repeat fold. Together, our work provides a flexible approach to detecting protein repeats and shows how their systematic annotation can reveal new aspects of protein diversity and evolution.

## Introduction

Protein domains arising from the repetition of specific repeat units are widespread across all domains of life, giving rise to a broad spectrum of symmetric topologies (1). Repetition has been, and continues to be, a major source of molecular innovation, producing ubiquitous protein folds such as TIM barrels, β-trefoils, and β-propellers (2–4) with the latter still emerging through ongoing blade amplification (4, 5). Owing to their modular organization, repeat proteins have long served as model systems for studying protein structure, function, and evolution, and as versatile scaffolds for protein engineering (6–8). Furthermore, their repetitive nature provides a powerful biophysical window into protein folding and enables systematic dissection of repeat-by-repeat energetics and cooperativity (9).

Across these diverse contexts, understanding repeat proteins requires the precise identification of repetitive regions, including their boundaries, copy numbers, and arrangement along the sequence. In particular, accurate identification of repeat units is important for studying the evolution of repeat architectures, as these units are the building blocks whose amplification, diversification, and rearrangement drive this process. The need for such detailed annotations has also grown with the rapid expansion of protein sequence and structure resources. Recent large-scale studies have uncovered extensive previously uncharacterized structural diversity in the AlphaFold Database (10), including thousands of highly symmetric and repetitive architectures (11), while complementary sequence-based analyses have revealed widespread and highly diverged tandem repeats across the tree of life (12). Consequently, developing accurate and scalable methods for repeat analysis remains a key task in protein science.

Various bioinformatics tools have been developed for the detection and analysis of protein repeats, ranging from early single-sequence approaches (13–15) that infer repeats from sequence patterns, periodicity, and recurring sequence motifs, to self-alignment (SSA) approaches in which a query sequence is aligned against itself to identify off-diagonal high-scoring regions indicative of repetitive patterns (16, 17). However, these methods are challenged by the divergence of repeat unit sequences, making their similarity difficult to detect (18). To overcome this limitation, modern repeat detection methods do not analyze the query sequence alone, but also exploit its homologs to amplify subtle similarity signals (18). For example, the early SSA implementations operating on a single sequence were later extended to incorporate homology information, resulting in HHrepID (19), one of the most sensitive tools, in which homologous sequences are first collected to construct a multiple sequence alignment (MSA), enabling sensitive profile-profile self-comparison. In contrast to SSA methods, profile-library-based methods incorporate evolutionary information in curated profiles of known repeat families. These precomputed profiles are then used to scan query sequences for regions matching specific repeat types, as implemented in the general HMMER/Pfam framework (20, 21) and in specialized methods such as REP/REP2 (22, 23) and TPRpred (24). Despite their algorithmic differences, both strategies share a common principle: leveraging homology to enhance similarity signals that might otherwise remain undetected in individual sequences.

Although the use of homology information has enabled detection of highly diverged repeats, its explicit incorporation also introduces practical limitations. In particular, the MSA construction required by sensitive alignment-based SSA methods is computationally expensive, limiting their application to large-scale studies. Profile-library-based methods are more tractable, but their sensitivity can be limited, and their dependence on manually curated, family-specific alignments reduces their flexibility. This motivates the use of representations that capture evolutionary and structural information relevant to repeat annotation without requiring explicit alignments. Recent advances in protein language models (pLMs) offer a promising framework for this problem by learning context-dependent sequence representations directly from unaligned sequences (25, 26). Trained on millions of sequences, pLMs provide residue-level representations (embeddings) that have emerged as an effective alternative to explicit MSA-based representations across tasks from structure prediction (27) to function annotation (28), largely because they encode information relevant to protein structure and evolution (29) and can be compared using simple similarity measures, as shown by nearest-neighbor approaches such as KnnProtT5 (30). Beyond direct similarity, these embeddings can be further refined through fine-tuning (31), contrastive learning (32), or embedding optimization (33), and have also been integrated into alignment-based frameworks, for example via context-dependent substitution matrices or differentiable end-to-end alignment models (34, 35).

High-sensitivity repeat domain detection relies on identifying divergent repeat units regardless of whether they are homologous or have arisen through convergent evolution. With this in mind, the ability of protein embeddings to capture evolutionary and structural information makes them natural candidates for this task. Building on this intuition, we previously developed pLM-Repeat (36), which performs self-alignment directly in ProtT5 embedding space to identify internal repetitive patterns and achieves comparable sensitivity to HHrepID without requiring time-consuming MSA construction. However, pLM-Repeat remains a *de novo* detection approach and does not address whether the prior-knowledge-driven paradigm of profile-library-based methods can be formulated within an embedding-based framework. Specifically, it remains unclear whether repeat annotation and detection can be cast as a supervised learning problem, and whether such a formulation could offer advantages.

In this study, we introduce TREAD (**<u>T</u>**ransfer learning-based **<u>RE</u>**peat **<u>A</u>**nnotation using Protein Embe**<u>D</u>**dings), a supervised framework for repeat annotation that leverages protein language model representations together with a deep neural network architecture trained on curated repeat datasets from RepeatsDB (8) or Pfam (21). These resources provide complementary forms of supervision. RepeatsDB supplies structurally defined repeat-unit annotations, whereas Pfam represents homologous repeat families through sequence alignments. Conceptually, TREAD resembles classical profile-based methods, but replaces MSA-derived profiles with trainable neural network models operating in embedding space. However, unlike these methods, TREAD does not necessarily assume homology between repeat units, as it can also be trained on structurally defined repeat classes. As a result, detecting the same repeat type in different proteins does not necessarily imply homology between them. TREAD provides a flexible alternative to profile construction, enabling residue-level scoring, motif boundary delineation, and large-scale annotation. Through our systematic benchmark against HMMER, a widely used tool for profile-based protein annotation, we show that the embedding-based supervised formulation often outperforms the profile-based approach, especially in low-data settings and when annotating divergent repeat families. We further demonstrate the practical utility of TREAD by applying it to a comprehensive analysis of β-propeller proteins, enabling the discovery and characterization of β-propeller diversity at scale.

## Materials and methods

### Model architecture and training

The TREAD model is an embedding-based neural network designed for residue-level repeat classification and scoring (Figure 1A). It integrates local convolutional feature extraction with long-range contextual modeling to generate residue-wise repeat score profiles. The model takes as input a per-residue embedding *X* ∈ R *^L×d^* extracted from protein language models, where *L* and *d* denote the sequence length and the embedding dimension (1024 in this study using the ProtT5 model, https://huggingface.co/Rostlab/prot_t5_xl_uniref50), respectively. ProtT5 was selected because it provides contextual residue-level representations and had previously shown utility for detecting distant protein relationships, including repetitive patterns in our previous pLM-Repeat study (35–37). To capture short-range sequence patterns, we applied a one-dimensional convolutional layer (Conv1D), followed by one or more residual convolutional (ResNet) blocks. Each block consists of two Conv1D-BatchNorm-ReLU-Dropout layers. To incorporate long-range contextual information, the convolutional features are subsequently passed to a single-layer bidirectional LSTM. The final prediction head comprises two fully connected layers with dropout, producing one or more task-specific scalar scores for each residue, depending on the prediction task.

**Figure 1.**
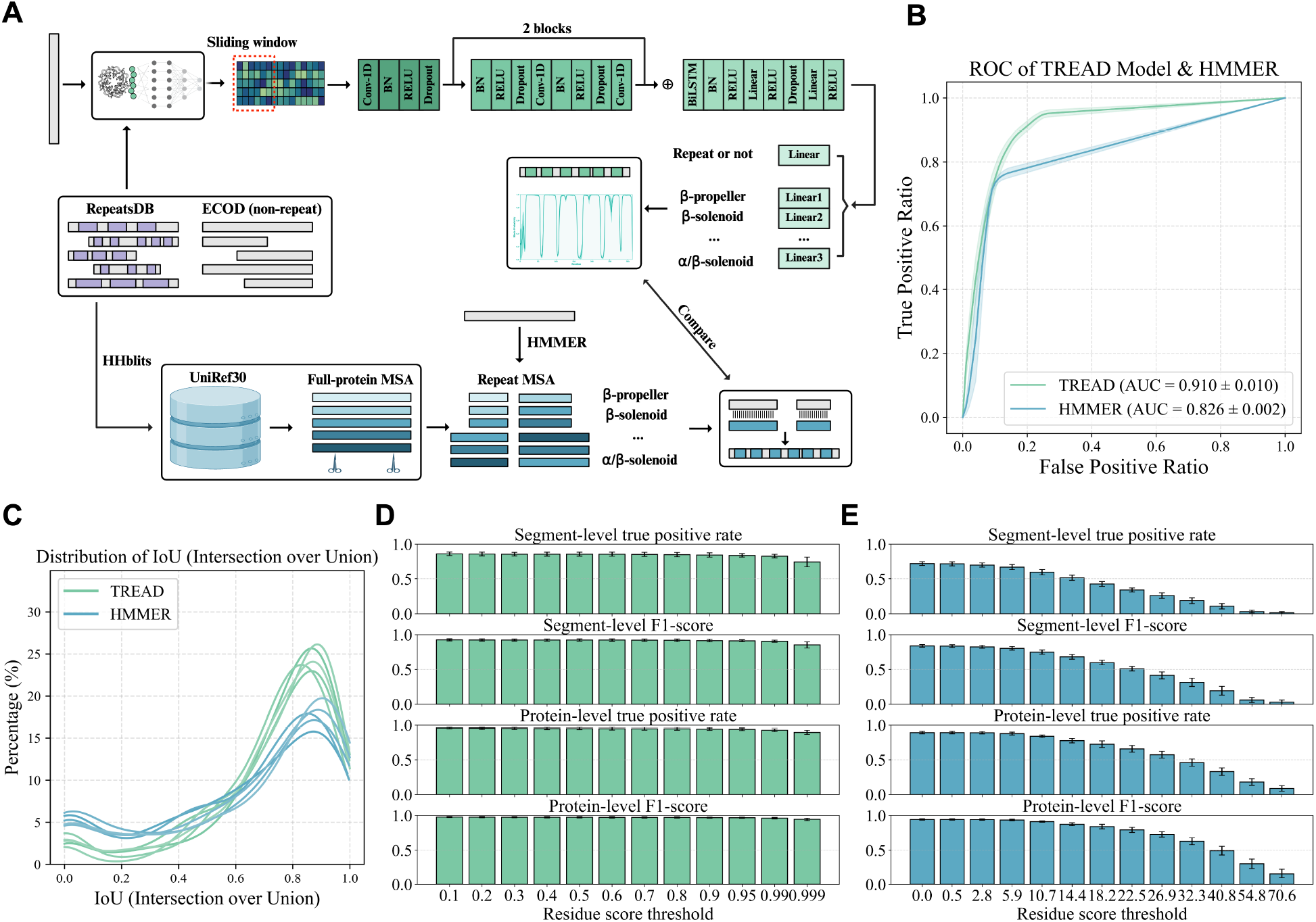
RepeatsDB benchmark design and performance comparison. (A) Overview of the benchmarking workflow for constructing the TREAD model and the HMMER pipeline. TREAD was trained on a dataset of RepeatsDB-annotated repeat proteins with a negative set sampled from ECOD. In contrast, the MSAs required by HMMER were generated by HHblits searches against UniRef30. The TREAD model consists of seven independent prediction heads, corresponding to six repeat fold classes and an additional binary head for repeat versus non-repeat prediction (see Materials and Methods for details). (B) Residue-level receiver operating characteristic (ROC) curve of TREAD (AUC = 0.910) and HMMER (AUC = 0.826) on the test dataset, shown in green and blue, respectively. Mean ROC curves across five cross-validation folds are shown as solid lines, with shaded regions indicating ±1 standard deviation. (C) Distributions of Intersection over Union (IoU) scores at the most permissive score thresholds (TREAD residue score threshold = 0.1; HMMER bitscore threshold = 0.0). Each method is represented by five curves corresponding to independent cross-validation folds. Only IoU scores from positive repeat sequences are included. (D) Segment-level true positive rate and F1-score of TREAD across a range of residue score thresholds at the minimum repeat length of 20. Bar heights indicate mean performance over five cross-validation folds, and error bars denote the variability across folds. (E) Corresponding segment-level performance of HMMER evaluated under the same protocol as in (D).

During training, fixed-length sequence windows (*W* = 64) were generated by partitioning each protein sequence into overlapping segments with a stride of 32 residues and a small random shift (±4 residues) applied to all window start positions to avoid placing window boundaries at the same positions in every training pass. The training windows inherited the corresponding residue-level labels for annotated repeat units from the full-length annotated sequence, as described in the following sections. To facilitate the extraction of contiguous high-scoring segments from per-residue scoring profiles returned by the model, we employed a soft-edge labeling scheme during training that promotes peak-like score profiles over annotated repeat units. Residues located in the repeat unit core were assigned a target label value of 1, whereas residues within the first and last 10% of the annotated repeat unit were linearly down-weighted to a minimum value of 0.5. Gaussian noise was added to the sequence embeddings during training as a regularization strategy to reduce overfitting. Finally, during inference, non-overlapping windows of length 64 were used to scan the full-length sequence to produce residue-level prediction profiles. We also tested partially overlapping windows with score averaging, but this did not improve performance. For the ablation study, we either replaced ProtT5 embeddings with one-hot embeddings or retained only the final linear layer of the TREAD architecture to produce residue-level scores.

### RepeatsDB benchmark scenario

We first designed a benchmark based on the RepeatsDB database to compare the performance of TREAD and HMMER. Using RepeatsDB v4.1 (https://repeatsdb.bio.unipd.it/) (8), we selected six abundant repeat protein folds: all-α solenoid, all-β solenoid, β-α solenoid, TIM barrel, β-barrel, and β-propeller. All RepeatsDB-PDB entries belonging to these fold classes were collected and clustered using MMseqs2 (version 89f75fec7fdbb0f3be99801a7b2b75e8f2895c5f, https://github.com/soedinglab/MMseqs2) with a maximum sequence identity of 90% and a minimum coverage of 80% (38), resulting in 1,823 repeat proteins annotated at the residue level for repeat unit boundaries and assigned to one of six fold classes. In addition, we sampled an equal number of non-repetitive sequences from the negative dataset of our previous study as non-repeat sequences (36), yielding a dataset of 3,646 proteins with residue-level annotations in total. Performance was assessed using five-fold cross-validation (CV) with one fold used for testing and the remaining four for training. The splitting of positive and negative examples into testing and training sets was guided by the MMseqs2 clustering results to ensure that the maximum sequence identity between the training and test sets did not exceed 30%, thereby minimizing homologous leakage.

Prior to the TREAD model training, hyper-parameter optimization was performed via grid search using the training set of one CV split, which was further randomly divided into training and validation subsets at an 80:20 ratio. Hyper-parameters were selected based on the residue-level F1-score on the validation set (SI Figure 1; SI Table 1). After hyperparameters were selected (64 output channels per residue, 2 ResNet blocks, kernel sizes of 3 for the initial convolutional layer and 7 for the ResNet blocks, and a hidden dimension of 64 in the penultimate fully connected layer), five independent models were trained, one for each CV split. Model training was carried out using the Adam optimizer with a learning rate of 1×10^-4^ in PyTorch (version 2.5.0), and early stopping with a patience of 10 epochs was applied. During evaluation, sequences in each test set were passed through the corresponding trained model to obtain residue-level prediction scores.

For the HMMER baseline, MSAs were constructed for the positive RepeatsDB-PDB entries in the training folds of each CV split by searching their sequences against the UniRef30 database (version 2023-02, https://colabfold.mmseqs.com/) using HHBlits (version 3.3.0, https://github.com/soedinglab/hh-suite) with default settings (39, 40). Although this procedure may introduce some information leakage through homologs related to positive test sequences, any such effect would be expected to favor HMMER rather than TREAD. The resulting full-length alignments corresponding to the positive samples were cut into repeat-unit alignments according to the annotated repeat boundaries. These repeat alignments were then converted into HMM profiles and assembled into a searchable profile database in the HMMER format using the *hmmpress* command from HMMER3 (version 3.2, https://hmmer.org/) (20). The same five CV splits used for TREAD were also used to construct and evaluate the HMMER baseline. For each CV split, sequences in the test set were searched against the associated HMM database using the *hmmscan* command with a highly permissive E-value threshold of 10000 to retain all potential hits for downstream thresholding analysis.

### Postprocessing of annotation results and evaluation

To ensure a fair and directly comparable evaluation of TREAD and HMMER, both methods were assessed using identical CV splits and an identical postprocessing and analysis framework. For each test sequence, TREAD produced residue-wise binary repeat/non-repeat scores alongside six fold-specific scores for the six repeat folds modeled by TREAD. For repeat sequences, all residues were scored using the prediction head corresponding to the ground truth fold. For non-repeat sequences, residues were scored using the binary prediction head. In parallel, HMMER outputs were parsed and projected onto the residue level by assigning each residue the bitscore of the segment to which it belongs. If a residue was covered by multiple HMMER hit segments, the maximum bitscore was retained. Similarly, only HMMER hits from the ground-truth fold were used for score assignment. Although HMMER is not natively designed to produce residue-level scores, this minimal projection retains the original segment-level scores while enabling boundary-aware comparison with the residue-level predictions produced by TREAD. Once the outputs of both methods were available at the residue level, performance was evaluated at the per-residue and per-segment levels.

### Residue-level evaluation

Each residue has a binary ground-truth label indicating whether it belongs to any of the six targeted repeat types. To ensure score availability for all residues, residues not covered by any HMMER hit were assigned a bitscore of *min-4* as the predicted score (*min* is the minimum bitscore reported across all hits in the benchmark). Under this formulation and using the score assignments defined above, residue-level receiver operating characteristic (ROC; Figure 1B) curves were computed for both TREAD and HMMER using scikit-learn (version 1.8.0, https://scikit-learn.org/stable/) (41).

### Segment-level evaluation

Predicted repeat segments were obtained by scanning residue-wise score profiles defined above and extracting all contiguous regions in which scores exceeded a predefined threshold *S*, and only segments with length ≥ *L* were retained. The predicted segments can correspond to the individual repeat units, but may also match only partially or cover more than one repeat unit. To assess the agreement between the predicted segments and ground-truth repeat units, the Intersection over Union (IoU) metric was used, with annotated repeat units and predicted repeat segments represented as intervals along the sequence (Figure 1C). For each protein in the test set of a given CV fold, IoU was defined as the ratio between the number of residues shared by annotated and predicted intervals and the total number of residues covered by either. In the subsequent evaluations (Figure 1D–E), a repeat protein from the test set was counted as a true positive if the IoU between annotated and predicted repeats exceeded 0.5, and as a false negative otherwise. Meanwhile, negative sequences were considered false positives if repeat segments were predicted. In addition to this strict IoU-based criterion, a relaxed presence/absence criterion was also applied in which a positive sequence was counted as a true positive if at least one repeat segment was predicted, whereas a negative sequence was counted as a true negative if no segment was predicted. In both the strict and the relaxed scenarios, sensitivity (true positive rate) was calculated from true positive and false negative counts, whereas F1 score was calculated using true positive, false negative, and false positive counts.

### Score thresholding

TREAD outputs a continuous score (or scores, depending on the setup) in [0,1] for each residue, enabling straightforward segmentation using a fixed set of score cutoffs [0.1, 0.2, 0.3, 0.4, 0.5, 0.6, 0.7, 0.8, 0.9, 0.95, 0.99, 0.999]. In contrast, absolute bitscore values in HMMER depend strongly on profile composition and database statistics, making fixed thresholds inappropriate. We therefore defined bitscore cutoffs using empirical quantiles of the residue-level bitscore distribution to obtain operationally comparable threshold ranges for segment extraction. For each benchmark, thresholds were set at quantiles [0.1%, 1%, 5%, 10%, 20%, 30%, 40%, 50%, 60%, 70%, 80%, 90%], computed over all bitscores, including the assigned min-4 values for residues without HMMER hits, and used for segment extraction. This resulted in threshold sets of [0.5, 2.8, 5.9, 10.7, 14.4, 18.2, 22.5, 26.9, 32.3, 40.8, 54.8, 70.6] for the RepeatsDB benchmark and [13.1, 17.3, 22.0, 24.7, 28.2, 31.0, 33.8, 36.7, 39.9, 43.6, 48.4, 55.4] for the Pfam benchmark (see below). An additional threshold at 0.0 was included to represent the most permissive setting.

### RepeatsDB TREAD model scan on AFDB Swiss-Prot

To further validate the generalization of the TREAD RepeatsDB model, we evaluated the averaged predictions of the five CV models using AFDB entries curated in the recent RepeatsDB update (8), with annotations generated using STRPsearch (42). The five models were applied to the same AFDB Swiss-Prot dataset (version 4, https://alphafold.ebi.ac.uk/download) used in the STRPsearch-based RepeatsDB update (8). Residue-level prediction scores were averaged for all entries, and repeat segments were extracted by identifying the contiguous high-scoring regions under different parameter combinations of *S* and *L* (Figure 2). For additional de novo predictions, repeat fold type was assigned using the fold-specific output with the highest residue-level scores. For each additional predicted entry, the predicted segments were extracted from the corresponding AlphaFold Model and compared pairwise using TM-align (43).

**Figure 2.**
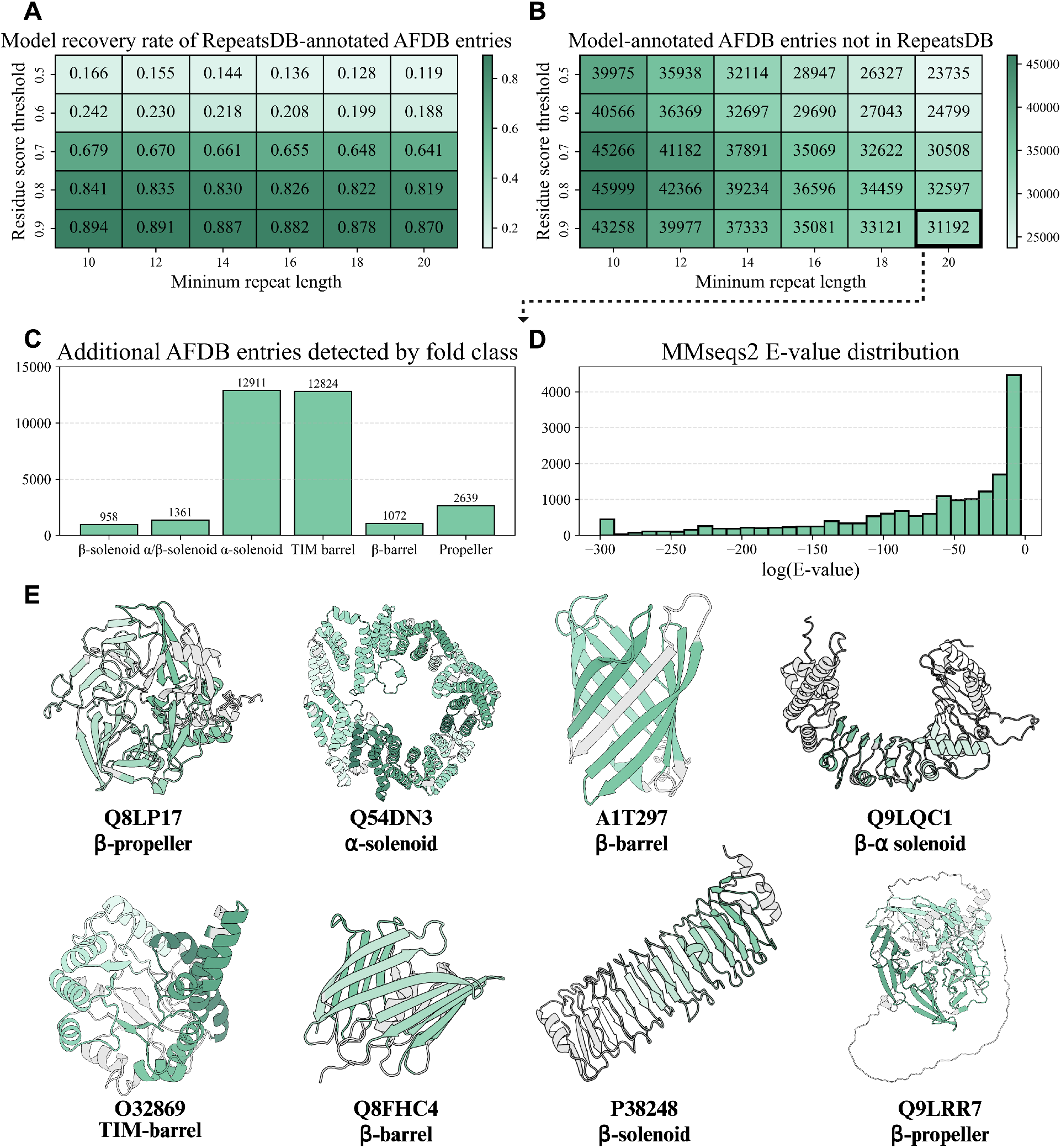
Swiss-Prot scan using the TREAD model. (A) Recovery rate on RepeatsDB-annotated proteins within AFDB, evaluated using the TREAD model. Results are summarized as a heatmap across two segment extraction parameters: the minimum residue-level score threshold (*S*) and the minimum repeat segment length (*L*). (B) Number of additional AFDB proteins predicted to contain repeats by TREAD but not annotated in RepeatsDB, shown across the same parameter grid as in (A). (C) Fold-class composition of the additional proteins detected by TREAD under a representative parameter setting (*S* = 0.9, *L* = 20). (D) Distribution of the best (minimum) MMseqs2 E-values obtained by searching the additional detected proteins against the RepeatsDB-AFDB set. Approximately 41% of these proteins show a significant match with an E-value ≤ 1×10^−3^. (E) Representative TREAD predictions missed by STRPSearch and lacking significant MMseqs2 similarity to annotated RepeatsDB-AFDB entries (E-value > 1×10^−3^), illustrating diverse repeat fold architectures considered in the model training.

### Evaluation of additional *de novo* repeat detectors in the RepeatsDB benchmark scenario

To provide a complementary comparison with *de novo* repeat-detection methods, RADAR (version 1.3, https://github.com/AndreasHeger/radar) (16), TRUST (version 1.0, https://zeus.few.vu.nl/programs/trustwww/) (17), HHrepID (version 1.1.0, https://toolkit.tuebingen.mpg.de/tools/hhrepid) (19), pLM-Repeat (version 1.0, https://github.com/KYQiu21/plmrepeat) (36), and the structure-based method CE-Symm (version 2.2.2, https://github.com/rcsb/symmetry/tree/master) were evaluated on the RepeatsDB benchmark using their default thresholds (44): RADAR repeat-family score = 100, TRUST P-value = 0.01, HHrepID P-value = 0.001, pLM-Repeat self-alignment score = 0.3, and CE-Symm self-alignment score = 0.4. IoU distributions were calculated as described above. As these methods do not require task-specific training data, they were evaluated once on the RepeatsDB entries pooled across the five TREAD/HMMER test folds. pLM-Repeat, the best-performing *de novo* sequence-based method in this comparison, was additionally evaluated at score thresholds of 0.1, 0.2, 0.25, 0.3, 0.35, 0.4, and 0.45.

### Pfam benchmark scenario

The second benchmark scenario utilized curated repeat alignments from the Pfam database (version 37.4, https://www.ebi.ac.uk/interpro/) (21). Eight repeat-unit families with sufficiently large Pfam full alignments were considered: WD40 (PF00400), KELCH (PF01344), ANK (PF00023), ARM (PF00514), HEAT (PF02985), TPR (PF00515), LRR (PF00560), and RCC (PF00415). After excluding sequences shorter than 70% of the family mean, the remaining sequences in each repeat family were partitioned into five folds using the same cross-validation procedure as in the RepeatsDB benchmark scenario; in each iteration, one fold was used for testing and the remaining four for training, with dataset splitting guided by MMseqs2 clustering to ensure that sequence identity between the training and test sets did not exceed 30%. For computational efficiency, if a split exceeded 8,000 training sequences or 2,000 test sequences, a random subsampling was performed to cap the datasets at these respective sizes. For each cross-validation split, negative sequences were independently sampled for the training and test sets using the same MMseqs2 clustering procedure, with the number of negatives matched to the corresponding number of positive sequences in each set. All negative samples were drawn from the ECOD30 database, excluding entries belonging to the X-groups Repetitive alpha hairpins, beta-propeller-like, and single-stranded right-handed beta-helix.

These compiled datasets served for both TREAD model training and HMMER profile construction. First, family-specific TREAD models for each repeat type were trained using the same hyperparameters as in the RepeatsDB benchmark. Second, for comparison with HMMER, repeat unit profiles were generated from the positive training sequences of each repeat family in each split and the resulting profiles were subsequently used to search the corresponding test set sequences and collect hits for evaluation. Finally, the evaluation procedure used in the RepeatsDB benchmark was applied here as well.

### Additional test sets in the Pfam benchmark scenario

Given that Pfam full alignments are generated with the *hmmsearch* tool of the HMMER package, the evaluation using Pfam alignments can be biased in favor of HMMER. To enable a more reliable assessment, we constructed two additional test datasets for the Pfam-based benchmark scenario:

### PSI-BLAST–derived test set

To reduce the potential bias, we used Pfam seed alignment of each repeat type as a query to search the nr30 database (version 4, https://www.ncbi.nlm.nih.gov/) using PSI-BLAST (version 2.13.0, https://blast.ncbi.nlm.nih.gov/Blast.cgi), with an E-value threshold of 1×10^-3^ and five iterations (45). For each repeat type, all retrieved hits were clustered using MMseqs2 with a maximum sequence identity threshold of 30% and a minimum coverage of 80%, resulting in the test dataset used to evaluate both methods. Each sequence was evaluated using all five cross-validation models trained for that repeat type, and the residue-level scores were averaged for final assessment. Similarly, residues were assigned a score by each five cross-validation HMM models and averaged for assessment.

### Pfam Clan–based remote homolog test set

To further evaluate the capability of both methods to detect remote homologs, we constructed additional datasets based on Pfam Clans. For each repeat type, other Pfam families from the same Clan were collected. Pfam families within the same Clan are considered homologous but too divergent to be represented by a single profile. From these remote homologous proteins, we randomly extracted at most 20,000 sequences to construct a remote homolog test set for each repeat type. Each sequence was evaluated using all five cross-validation models trained for that repeat type, and the residue-level scores were averaged for final assessment.

### Scale study using WD40

To investigate how training data availability influences the relative performance of the learning-based TREAD and profile-based HMMER approaches, we performed a data scaling experiment using the WD40 dataset from the Pfam benchmark as a representative case. The MMseqs2 clusters were randomly shuffled and partitioned into twenty folds to prevent homologous leakage. Nineteen folds were combined to form a fixed test pool, from which we sampled 20,000 WD40 sequences in a cluster-balanced manner, given the large size of the WD40 alignment. Similarly, 20,000 non-propeller sequences sampled from the ECOD30 database served as negative test samples.

From the remaining fold, we constructed nested training subsets by selecting the first fraction *p* of MMseqs2 clusters in the shuffled order, ensuring that larger subsets fully include all sequences from smaller subsets, with *p* ∈ [0.01, 0.02, 0.05, 0.1, 0.2, 0.5, 1.0]. For each *p*, all sequences within the selected MMseqs2 clusters formed the positive training set, while an equal number of negative sequences were sampled from the ECOD30 database (excluding those sampled previously for the test set) using the same nested strategy as for the positive training subsets. For robustness, we generated five independent realizations of these subsets. In each repetition, each subset at a given *p* was used for model training and profile construction (only positive sequences), and performance was evaluated on the fixed test set prepared as described above.

### Construction of the propeller blade model

To construct a propeller blade-specific TREAD model, all ECOD domains belonging to the beta-propeller-like X-group (ECOD ID: 5.1) were extracted and clustered using MMseqs2 with 80% sequence identity and 80% coverage. The resulting 781 propeller domains were manually inspected to annotate propeller blades, i.e., parallel four-stranded β-meanders (β1-β2-β3-β4) and label blade residues as positive. We further randomly sampled 200 ECOD domains from each ECOD Architecture, excluding the beta-propeller-like X-group, resulting in a total of 3,536 negative domains, in which all residues were assigned zero label, after clustering at 80% sequence identity. The combined dataset was clustered at 30% sequence identity using MMseqs2, followed by an 80/20 cluster-based split, ensuring no sequence identity above 30% between the training and test sets.

Hyperparameter optimization was performed via grid search using five-fold cross-validation on the training set (SI Figure 16). During training, 20% of the sequences in each training fold were held out as a validation set to monitor performance, with an early stopping criterion of 10 epochs applied to prevent overfitting. The optimal hyperparameter configuration, comprising 128 output channels per residue, a single ResNet block, kernel sizes of 11 for the initial convolutional layer and 7 for the ResNet blocks, and a hidden dimension of 64 in the penultimate fully connected layer, was selected based on the mean residue-level F1 score across the five folds (SI Table 3). The final model was trained on the full training dataset for a number of epochs corresponding to the average convergence epoch observed during the grid search.

### Propeller blade model scan on AFDB50

The propeller blade model was used to scan the AlphaFold Database (version 6, https://alphafold.ebi.ac.uk/download) clustered at 50% maximum sequence identity (AFDB50) (46, 47). Each sequence was converted into ProtT5 embeddings and passed through the model for prediction. Discrete blade regions were identified by segment extraction using a score threshold of 0.8 and a minimum length threshold of 20 residues.

Only entries containing at least three blades were retained. For each entry, the sequence was truncated to span the predicted propeller region, defined as the region from the start of the first detected blade to the end of the last detected blade. The resulting truncated sequences were filtered to retain those with an average pLDDT ≥ 75, hereafter referred to as UniPropeller. General metadata for all entries (structure, taxonomy, pLDDT, etc.) were retrieved via the AFDB API. In addition, putative functional annotations were assigned using eggNOG-mapper (version 2.1.13; https://github.com/eggnogdb/eggnog-mapper) in Diamond mode with an E-value cutoff of 1×10^-3^ (48).

For subsequent characterization, UniPropeller sequences were clustered at 50% sequence identity using MMseqs2, resulting in the UniPropeller50 set. We searched these sequences against all ECOD propellers using the MMseqs2 *easy-search* method with maximum sensitivity (-s 9.0), and assigned the propeller family of the best hit (E-value ≤ 1×10^-3^) to each sequence. For visual exploration of the sequence space, hierarchical clustering was performed using the ProteinClusterTool package (https://github.com/johnchen93/ProteinClusterTools) at the default settings (49), generating an interactive sequence map (Figure 5H). To further assess structural features, all UniPropeller50 proteins were searched against ECOD propellers using Foldseek with default parameters (version d60563224ad2cf976af580cc48b1568bd35b4716, https://github.com/steineggerlab/foldseek) (50). In addition, AFDB propeller domains curated in the TED database (https://ted.cathdb.info/) were retrieved for comparison with UniPropeller50 dataset (51). For comparison, we performed HMMER searches to collect homologs from the same AFDB50 dataset. Each propeller sequence used for model training was first used as a query to generate an MSA with HHblits against the UniRef30 database. The resulting alignments were then converted into HMM profiles, which were used to search AFDB50 using default settings. The majority of sequence analyses on selected cases were performed using the MPI bioinformatics toolkit (https://toolkit.tuebingen.mpg.de/) (52).

### Propeller blade model scan on representative proteomes

To obtain a distribution of propellers at a finer taxonomic scale, we further applied the blade model to a collection of representative proteomes spanning 4,618 species, curated in a recent study (12). Proteome sequence files were retrieved from NCBI (https://www.ncbi.nlm.nih.gov/refseq/), resulting in a total of 21,446,147 proteins. All sequences were scanned using the same pipeline as in the AFDB50 analysis, resulting in 283,783 sequences with at least three propeller blades detected. Putative functional annotations for all detected propeller-containing proteins were assigned using eggNOG-mapper, following the same protocol described above. The ggtree package (version 4.0.4, https://github.com/YuLab-SMU/ggtree) was used to visualize the phylogenetic tree shown in Figure 7A (53).

To further investigate evolutionary conservation of propeller architectures, we performed an orthology-based analysis along selected taxonomic paths. Orthologous protein pairs between species proteomes were obtained from the InParanoid database (version 9, https://inparanoidb.sbc.su.se/) (54). For each ortholog pair, proteins in which at least three propeller blades were detected in both species were classified as ortholog pairs with propellers detected in both proteins, suggestive of conserved propeller architectures.

### Visualization

Structures were visualized with PyMOL 3.0 (https://www.pymol.org/). Figures 1A, 3A, 4A and 5A were generated using BioRender.com. Other figures were plotted using the Matplotlib v3.10 and seaborn v0.13 packages (55, 56). Sequence alignments were visualized using ESPript 3.2 (https://espript.ibcp.fr/ESPript/ESPript/index.php) (57).

## Results

### TREAD model produces residue-level repeat predictions

TREAD is a sequence-based predictor that uses ProtT5 representations to generate residue-level repeat scores, from which contiguous segments, typically corresponding to repeat units, can be extracted (Figure 1A). Owing to the use of fixed-length sequence windows, the predictions are driven primarily by local sequence motifs rather than global sequence context, thus making the per-residue scores well suited for identifying repeat segments with defined boundaries. Depending on the setting, TREAD can be configured to support single or multi-type repeat annotation. This flexibility is exemplified in the benchmarks presented below.

### TREAD matches HMMER with less explicit evolutionary information

We compared TREAD with HMMER, a method widely used for the annotation of repeats based on matches to manually curated profiles of repeat families. To this end we designed a benchmark based on RepeatsDB (Figure 1A), which provides structurally curated residue-level annotations of repeat units. Six abundant repeat fold classes were considered: all-α solenoid, all-β solenoid, β-α solenoid, TIM barrel, β-barrel, and β-propeller. To assess whether ProtT5 representations combined with supervised learning can achieve performance comparable to that of HMM profiles derived from MSAs, which explicitly capture evolutionary information, TREAD was trained exclusively on RepeatsDB-annotated sequences, whereas HMMER relied on repeat-unit profiles constructed from these sequences and their homologs. Both methods were evaluated using five-fold cross-validation (CV) with a stringent sequence identity cutoff (<30%) between training and test sets (see Materials and Methods for details).

At the residue level, TREAD substantially outperformed HMMER, achieving an Area Under the Curve (AUC) score of 0.910±0.010 compared with 0.826±0.002 across five CV folds (Figure 1B). We next examined whether this residue-wise advantage translates into improved segment recovery. Predicted repeat segments were defined as contiguous regions exceeding a minimum length *L* = 20 in which residue scores surpassed a confidence threshold *S* (0.1 for TREAD and 0.0 for HMMER; see Methods for thresholding details), and evaluated against ground-truth annotations using the Intersection-over-Union (IoU) metric, with the IoU distribution shown in Figure 1C for repeat proteins in the test set only. Under these settings, TREAD yielded higher per-protein IoU values in each of the five CV folds (paired Wilcoxon signed-rank test, p < 0.0001 in all folds) and the same relative advantage of TREAD over HMMER was observed across score (*S*) thresholds (SI Figure 2). To assess segment recovery in a classification framework, we used an *IoU* threshold of 0.5 to define true positives, false positives, and false negatives (see Methods for details), and found that TREAD achieved the highest segment-level performance across the tested score thresholds, with peak F1 score and the corresponding true positive rate of 0.926 ± 0.013 and 0.863 ± 0.023, respectively, compared with 0.838 ± 0.017 and 0.721 ± 0.025 for HMMER (Figures 1D and 1E; two top panels). The same pattern was observed when classification was evaluated using a more relaxed presence/absence criterion: both methods achieved high performance, but TREAD retained a modest advantage. Across tested thresholds, TREAD reached higher peak F1 scores (0.977 ± 0.006 vs. 0.942 ± 0.009) and true-positive rates (0.956 ± 0.012 vs. 0.890 ± 0.015) than HMMER (Figures 1D and 1E; two bottom panels). Varying the minimum segment length *L* produced similar results under both the strict IoU-based and the relaxed presence/absence criteria, indicating comparable robustness to the minimal segment length selection (SI Figure 3; SI Figure 4). Notably, in the above benchmarks TREAD exhibited strong robustness to score threshold variation, whereas HMMER was highly sensitive to the choice of *S* with metrics improving substantially under relaxed thresholds. This robustness stems from the sharply bimodal residue-score distributions produced by the model, in which positive residues cluster near 1.0 and negatives near 0.0 (SI Figure 5), yielding a large decision margin that minimizes sensitivity to threshold selection.

**Figure 3.**
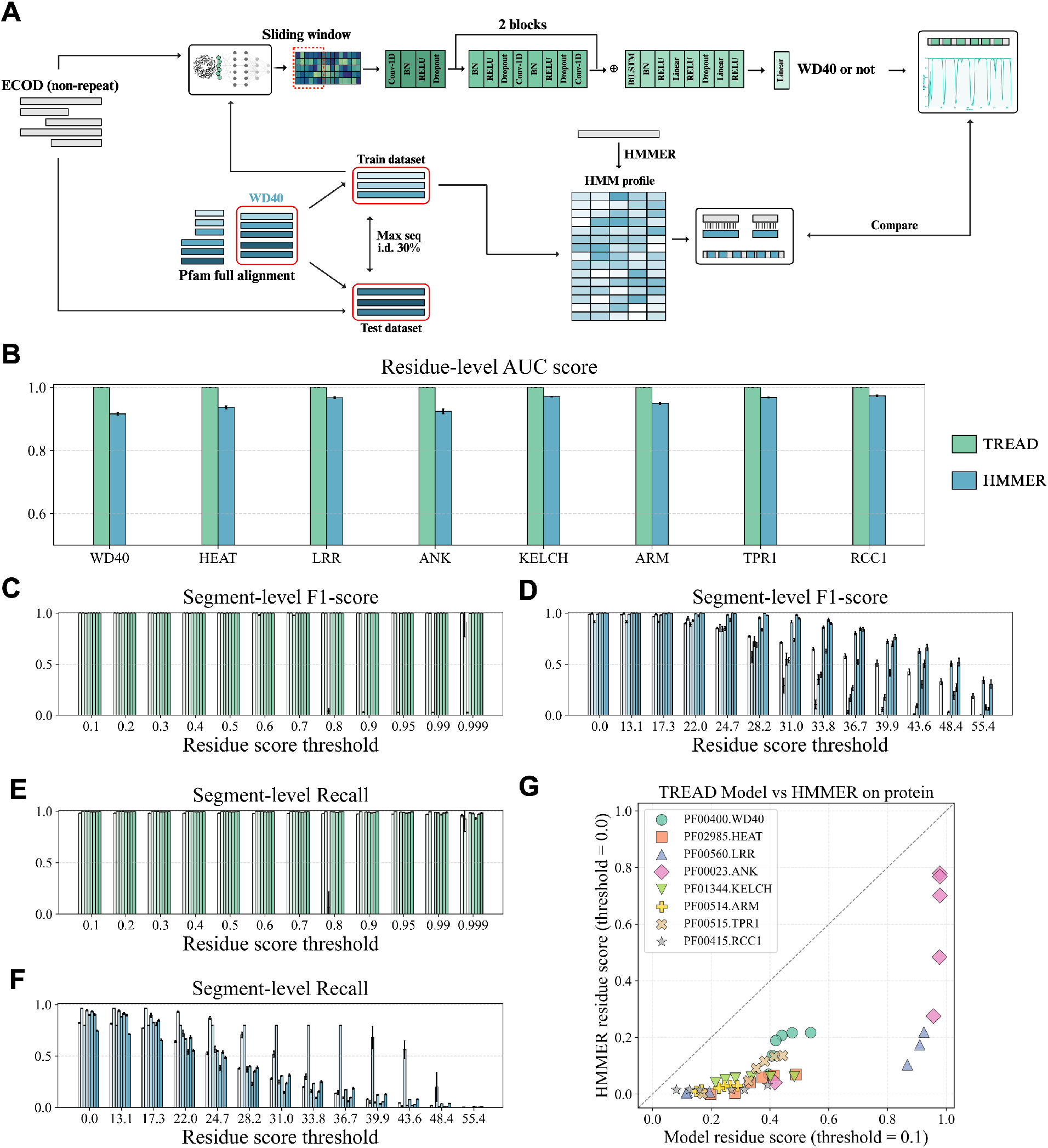
Pfam benchmark design and performance comparison. (A) Overview of the Pfam-based benchmarking protocol. For each repeat family, identical Pfam full alignments were used either to train a family-specific TREAD model or to construct the corresponding HMMER profile, ensuring a controlled and fair comparison. Eight repeat families were evaluated: WD40 (PF00400), HEAT (PF02985), LRR (PF00560), ANK (PF00023), KELCH (PF01344), ARM (PF00514), TPR1 (PF00515), and RCC1 (PF00415). Each TREAD model contains a single prediction head dedicated to one repeat family (see Materials and Methods). (B) Residue-level ROC curves for TREAD and HMMER across the eight Pfam families, evaluated on the test datasets derived from Pfam full alignments. Plotting conventions are identical to Fig. 1B. (C-D) Segment-level F1-scroe across residue score threshold at the minimum repeat length of 20 for (C) TREAD and (D) HMMER, evaluated on the Pfam full-alignment test datasets. At each threshold, bars (from left to right) correspond to WD40, HEAT, LRR, ANK, KELCH, ARM, TPR1, and RCC1. Error bars indicate variability across five cross-validation folds, following the same convention as Fig. 1D. (E-F) Segment-level Recall across residue score thresholds at the minimum repeat length of 20 for (E) TREAD and (F) HMMER, evaluated independent test datasets generated by PSI-BLAST. Bars are ordered and plotted identically to those in (C–D). (G) Recall comparison under the relaxed presence/absence criterion between TREAD and HMMER on datasets curated from Pfam families belonging to the same Pfam Clan for each repeat type. Both methods were evaluated under the most permissive detection setting (TREAD residue score threshold = 0.1; HMMER bitscore threshold = 0.0). For each repeat family, multiple minimum segment length thresholds (*L* = 10, 15, 20, 25, 30, 35) were tested and shown as individual scatters, with families distinguished by unique marker shapes and colors.

**Figure 4.**
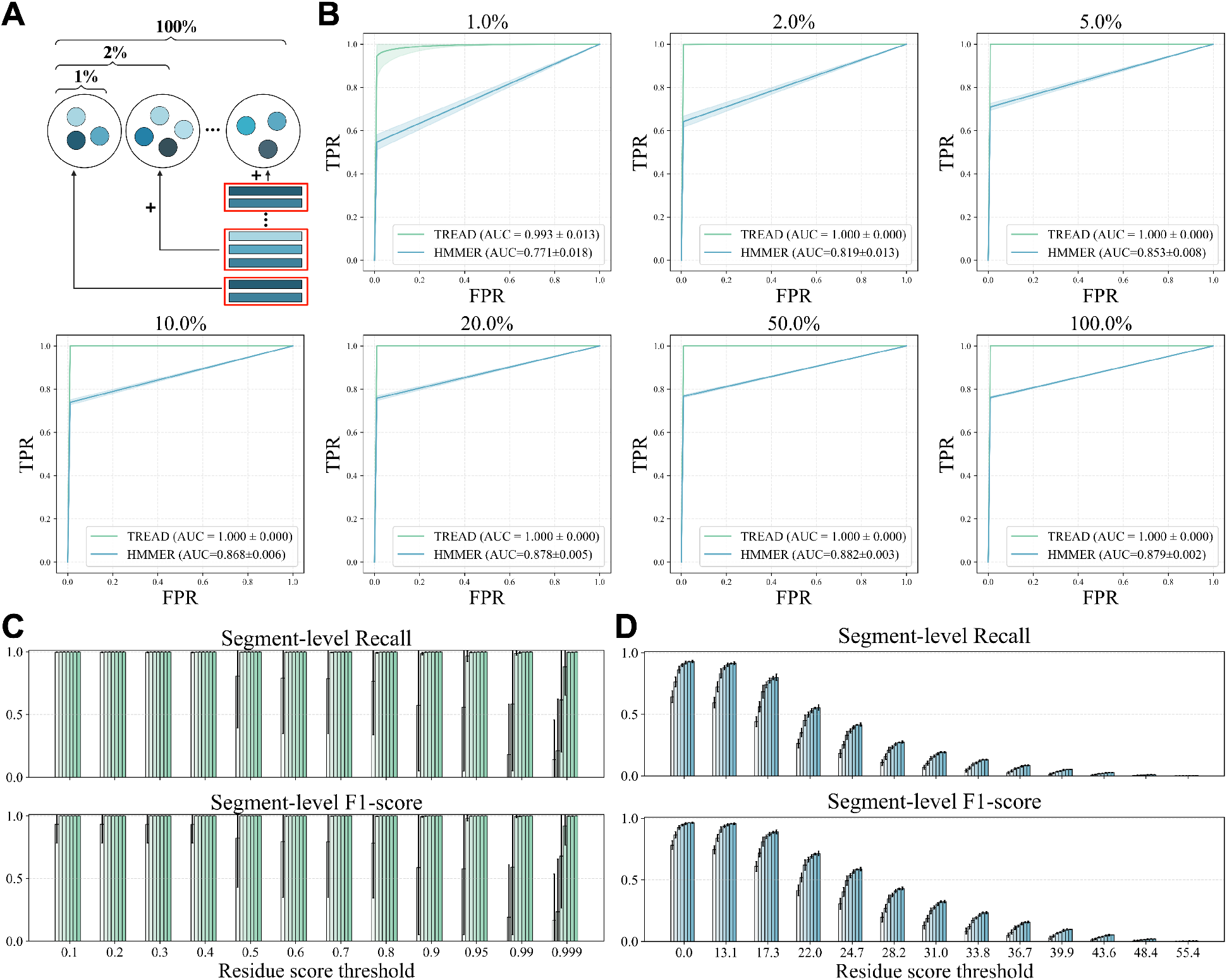
Scaling analysis of TREAD and HMMER performance on WD40. (A) Overview of the dataset construction design. A fraction *p* of the training dataset was used for model training or profile construction in a cluster-aware nested manner (*p* ∈ [1%, 2%, 5%, 10%, 20%, 50%, 100%]; see Materials and Methods for details). (B) Residue-level ROC curves for TREAD and HMMER under different training data fractions, with each panel corresponding to one value of *p*. Plotting conventions are identical to Fig. 1B. (C-D) Segment-level true positive rate and F1-score across residue score thresholds for (C) TREAD and (D) HMMER, evaluated under the same scaling settings as in (B) and plotted following the convention of Figure 3C-D.

To disentangle the contributions of input representation and model architecture, we performed an ablation analysis. Replacing pLM embeddings with one-hot encodings, or removing contextual modeling components while retaining only the final linear layer, led to substantial performance drops relative to the baseline (average residue-level F1 scores of 0.752 and 0.851 vs. 0.910, respectively; SI Table 2). These results underscore the importance of both informative protein representations and contextual modeling for accurate repeat annotation by TREAD.

To place the TREAD/HMMER comparison in the context of *de novo* repeat detection methods, we additionally evaluated RADAR (16), TRUST (17), HHrepID (19), pLM-Repeat (36), and the structure-based method CE-Symm (44) on the same RepeatsDB benchmark using their default settings (see Materials and Methods). Compared with TREAD and HMMER, these *de novo* approaches showed a substantially larger fraction of predictions at low or zero IoU and less pronounced enrichment at high IoU values, indicating poorer recovery of the curated repeat regions overall (SI Figure 6). Among the sequence-based *de novo* approaches, pLM-Repeat, which, like TREAD, relies on ProtT5 representations, showed the strongest performance, with a clear high-IoU peak despite operating without task-specific supervised training. We therefore further evaluated pLM-Repeat across a range of self-alignment score thresholds. More permissive thresholds improved repeat recovery, but even under the least stringent settings the IoU distribution remained clearly shifted below those of TREAD and HMMER (SI Figure 7). These results further support the advantage of prior knowledge-driven repeat annotation when the target repeat type is predefined.

In summary, TREAD achieves comparable or superior performance to HMMER on the RepeatsDB benchmark across residue-level and segment-level evaluations while using far less explicit evolutionary information, highlighting the strength of embedding-based models for repeat annotation.

### TREAD recovers RepeatsDB AFDB annotations and expands repeat discovery

To further assess the generalization of the TREAD model beyond the training data, we evaluated the averaged predictions of the five CV models on AFDB entries recently integrated into RepeatsDB, but not used for TREAD training. These domains were identified by STRPSearch (42), a structure-based repeat detection pipeline that uses Foldseek searches against curated structural repeat libraries (50). Although STRPSearch also follows a prior-knowledge-driven strategy, a direct head-to-head comparison with TREAD is not straightforward, as STRPSearch relies on manually curated structural databases and thus makes it difficult to construct a cleanly separated training and test dataset. Instead, we applied the five TREAD CV models to scan the same AFDB Swiss-Prot subset analyzed by STRPSearch to evaluate their ability to recover entries annotated as repetitive in RepeatsDB, providing a cross-method comparison on a shared sequence set.

Following criteria similar to those employed by STRPSearch, a repetitive domain was considered successfully recovered if at least three repeat segments were detected using the fold-specific output corresponding to the annotated fold type. Recovery rates, i.e., the fraction of repetitive domains that were successfully detected, were summarized as a heatmap across combinations of two segment extraction parameters: minimum segment length *L* and minimum residue-level score *S*. As shown in Figure 2A, increasing the score threshold *S* beyond 0.5 (corresponding to the minimum soft-label score for positive residues, see Materials and Methods) led to higher recovery rate while increasing stringency, likely because stricter thresholding yielded cleaner separation of individual repeat units. In contrast, restricting the minimum segment length *L* had only a minor effect on recovery. For example, under a representative parameter setting (*L* = 20, *S* = 0.9), TREAD successfully recovered 87.0% of the curated RepeatsDB-AFDB entries.

Notably, beyond recovering annotated domains, TREAD identified an additional set of proteins predicted, using the highest-scoring fold-specific output, to contain repeats from the six repeat fold classes but missed by STRPSearch, particularly enriched in α-solenoid and TIM-barrel folds (Figure 2B–C). A search using MMseqs2 against the annotated AFDB repeat set revealed that 41% of these predictions share significant sequence similarity with annotated repeats (E-value ≤ 1×10^-3^; Figure 2D), supporting them as plausible repeat-containing proteins that STRPSearch may fail to detect. Representative examples from the subset without significant MMseqs2 similarity to annotated AFDB repeats are shown in Figure 2E. To further assess whether these additional TREAD predictions missed by STRPsearch were supported at the structural level, we extracted the repeat segments and compared them pairwise within each protein using TM-align. The majority of intra-protein repeat-unit comparisons yielded TM-scores above 0.5 (SI Figure 8), indicating structural similarity among the regions independently identified from sequence-derived embeddings. This provides orthogonal structural support for many of the additional repeat predictions made by TREAD. Collectively, these results suggest that TREAD provides a complementary sequence-based approach for large-scale repeat discovery and database curation, further demonstrating the generalization ability of the proposed supervised learning framework.

### TREAD shows stronger generalization than HMMER on the Pfam benchmark

The strong performance of TREAD on RepeatsDB motivated a complementary evaluation using Pfam (Figure 3A), the largest sequence-based repeat protein resource. While RepeatsDB provides structural annotations for experimentally solved proteins, Pfam offers deep MSAs for many repeat families; however, unlike RepeatsDB entries, Pfam families correspond to individual repeat units rather than full repeat-containing proteins with annotated unit boundaries. We selected eight common repeat types from Pfam for this benchmark: WD40, KELCH, ANK, ARM, HEAT, TPR, LRR, and RCC. For each Pfam repeat unit family, the full alignment was partitioned into five folds, and in each fold both TREAD and HMMER were evaluated using exactly the same test sequences. Thus, unlike in the previous benchmark scenario, each Pfam family is associated with a dedicated TREAD model.

First, when evaluated on test sets derived from Pfam full alignments, both methods achieved near-ceiling performance across all repeat types. At the residue level (Figure 3B; SI Figure 9), TREAD exhibited almost perfect discrimination, with an average AUC of 1.0 across the eight families. HMMER also performed strongly, with AUC values consistently above 0.920. Segment-level evaluations also revealed similarly high accuracy for both approaches, with peak average F1-scores of 0.999±0.001 for TREAD and 0.986 ± 0.029 for HMMER across repeat types and CV folds (Figure 3C-D). Finally, across the extraction parameters *L* and *S*, both methods showed trends consistent with those observed in the RepeatsDB benchmark (SI Figure 10). The performance drop of TREAD observed for LRR (Figure 3C and E; third bar) at stringent score thresholds reflected its short repeat-unit length relative to the L = 20 cutoff used in these panels; however, this effect disappears at lower L values (SI Figure 10).

Since both methods performed nearly perfectly on test sets derived from Pfam full alignments, we considered two additional datasets to provide a more challenging assessment of generalization. These test sets were built to contain more diverse homologs that were less ideally matched to the family profiles represented in the Pfam full alignments. Additionally, because Pfam full alignments are constructed using HMMER, adding these extra datasets helped reduce a potential bias in favor of HMMER. First, we constructed a test set by using PSI-BLAST together with Pfam seed alignments to identify homologs in the NR30 database. Under this setting, the performance gap in the segment-level evaluations widened (Figure 3E-F; SI Figure 11): TREAD maintained high recall (0.989 ± 0.012), whereas HMMER performance declined significantly (Recall = 0.877 ± 0.074). This likely reflects the fact that a single HMMER profile cannot capture the broader sequence diversity present in the PSI-BLAST-derived test set. In addition, we evaluated both methods on remote homolog datasets derived from Pfam Clans. In Pfam, families within the same clan are considered homologous but too divergent to be represented by a single profile, and thus provide a potential set of remote homologs for each repeat type. Using relaxed presence/absence evaluation, TREAD consistently achieved higher recall (Figure 3G) and F1-score (SI Figure 12) than HMMER across a range of minimum segment length thresholds *L*. It should, however, be noted that the strongest apparent signal on the clan-based dataset was observed mainly for the ANK and LRR families.

Given the very good performance of TREAD in the Pfam benchmark and its variants, we also assessed its simplified version without any further contextual learning, i.e., with only the linear prediction head. The simplified model maintained excellent performance at the residue level on Pfam full alignments, achieving an average AUC score of 1.0 across all repeat types (SI Figure 13). On the PSI-BLAST–derived test set, both the full and simplified models achieved high peak recall (>0.950 across all repeat types), with a modest advantage for the full model (0.961 ± 0.058 for the linear model; SI Figure 13). Notably, in this benchmark the two models exhibited distinct behaviors under increasingly stringent score thresholds during segment extraction. Whereas the full TREAD model remained largely insensitive to threshold selection for most repeat types (Figure 3E), the simplified linear model showed a more gradual performance decline (SI Figure 14), resembling the behavior observed previously for HMMER. This difference suggests that contextual learning components stabilize residue-level confidence calibration. The benefit of architectural complexity became most pronounced on the Pfam Clan–based remote homolog dataset, where the simplified model consistently underperformed the full TREAD model across nearly all parameter settings (SI Figure 15).

Together, these results indicate that while TREAD and HMMER perform comparably when evaluated within information-rich Pfam MSAs, TREAD exhibits stronger generalization on datasets that deviate from the profile construction biases inherent in Pfam. The ablation analyses further suggest that the architectural complexity is not universally required, but becomes critical for robust repeat detection in more divergent and out-of-distribution scenarios.

### TREAD outperforms HMMER in the low-data regime

Given that both TREAD and HMMER depend on the availability of data for model training and profile construction, respectively, we next examined how their performance scales with the size of the training/profile-construction set. To this end, we performed a sample-efficiency analysis, in which only a fraction of the Pfam full alignment was used to train the model or construct HMM profiles. We performed this analysis using the WD40 family as a case study, given its large size and high sequence diversity. Specifically, we varied the fraction of sequences used for training from 1%, 2%, 5%, 10%, 20%, 50%, to 100%, using a clustering-based subsampling strategy to ensure that higher fractions strictly contained lower fractions (Figure 4A; Materials and Methods), and evaluated model performance.

At the per-residue level, TREAD exhibited strong robustness under reduced training data conditions. Even when trained on only 1% of the WD40 alignment (averaging 413 sequences per fold), TREAD achieved near-ceiling performance, with an average residue-level AUC of 0.993 across five independent realizations (Figure 4B). In contrast, HMMER showed a pronounced dependence on alignment size. With 1% of the alignment, HMMER achieved modest performance (AUC = 0.771 ± 0.018). Performance improved steadily as additional homologs were included, reaching AUC values of 0.853–0.868 at 5–10% of the alignment, and approaching its maximal range (0.878–0.882) when 20– 100% of the alignment was used (Figure 4B). A similar trend was observed at the segment level (Figure 4C-D): when trained on only 1% of the alignment data, TREAD and HMMER achieved peak F1-scores/Recall of 0.932 ± 0.133/0.998 ± 0.002 and 0.782 ± 0.031/0.643 ± 0.024, respectively, at the loosest score threshold (0.1 for TREAD and 0.0 for HMMER; minimum residue score threshold during segment extraction).

Taken together, these results reveal a difference in how the two approaches respond to limited training/profile-construction data. TREAD, likely aided by the use of pretrained ProtT5 embeddings that already encode substantial evolutionary information at the single-sequence level, was more sample-efficient in this benchmark, achieving stable performance from relatively small training sets, whereas HMMER must derive such evolutionary information from increasingly deep and diverse alignments for effective profile construction.

### Exhaustive AFDB50 scan characterizes a comprehensive and diverse β-propeller landscape

Encouraged by the robust benchmark performance, we next applied the TREAD framework to a large-scale case study focusing on β-propellers. As one of the largest and most diverse repeat protein families, β-propellers lack a general sequence-based detection method applicable across the entire protein fold, despite existing tools for specific subfamilies, such as WDSP for WD40 (58), and PropLec for propeller lectins (59). Using a curated dataset obtained from the ECOD database (see Methods for details) with propeller blade regions manually annotated, we trained a propeller-specific TREAD model that achieved a per-residue F1-score of 0.945 on an independent test set (Figure 5A). This model was subsequently used to scan the latest AFDB50 database (49,851,774 sequences) to identify propeller blade residues.

**Figure 5.**
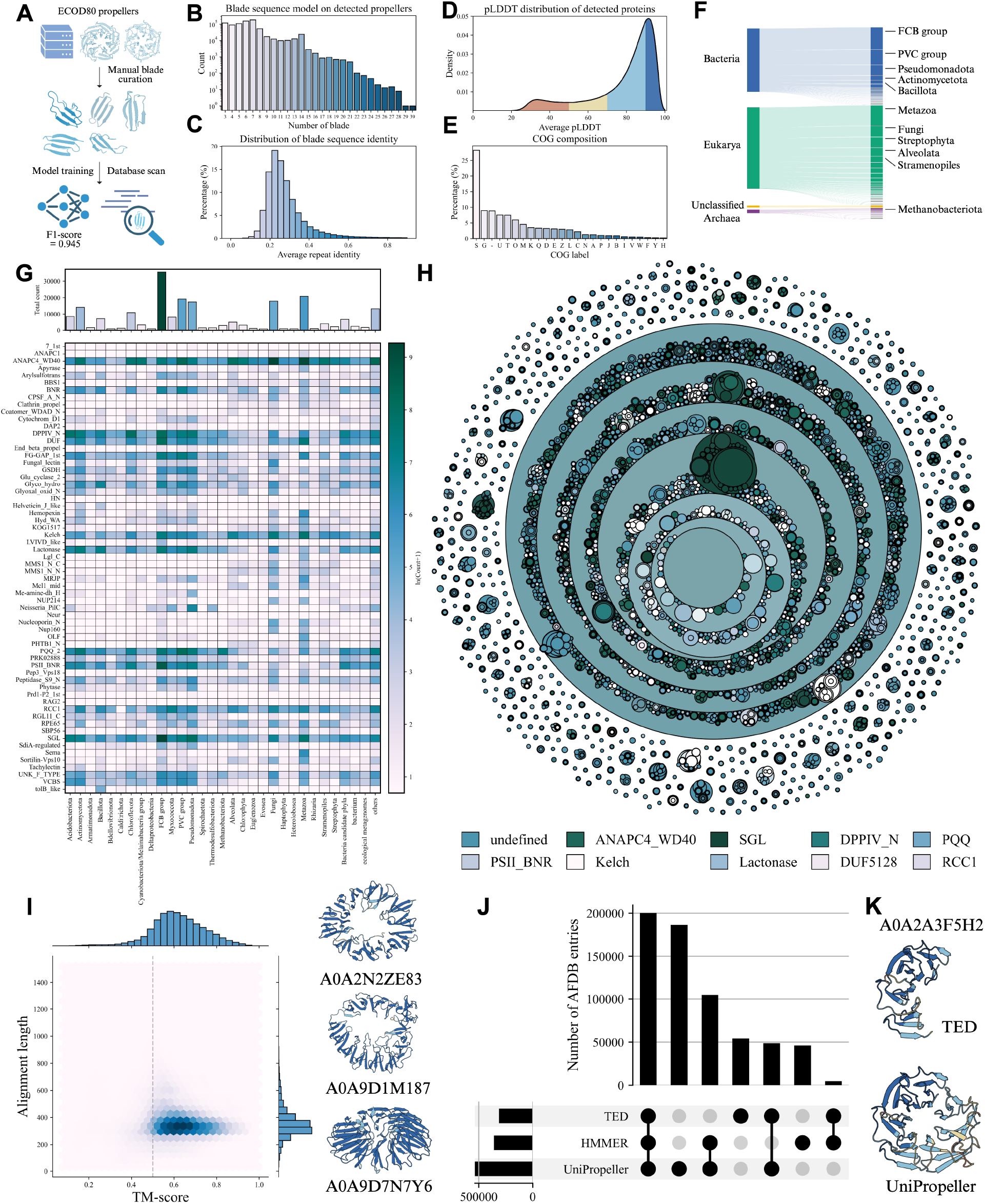
Propeller landscape obtained from the exhaustive AFDB50 scan. (A) Schematic overview of the construction and application of the TREAD propeller detection model. (B) Distribution of blade copy numbers among all detected entries containing at least three blades, shown on a logarithmic scale. (C) Distribution of blade sequence identity, where each entry is represented by the average pairwise Needleman–Wunsch alignment identity among all detected blades within the same protein. (D) Distribution of AlphaFold model pLDDT scores for all detected entries. For each entry, only the region between the first and last predicted blades was used to calculate the average pLDDT score. (E) COG functional category composition of all entries containing at least three blades (see Supplementary Table 4 for category definitions). (F) Taxonomic distribution of detected propeller-containing proteins across major phyla and super kingdoms. Numbers indicate the total counts of domains within sequence clusters assigned to each taxonomic level. (G) Heatmap of propeller family enrichment across phyla. To ensure data quality and computational efficiency, only entries with an average pLDDT ≥ 75 were retained and clustered at a maximum sequence identity of 50%. Remaining entries were assigned to ECOD propeller families (F-groups) using MMseqs2. The top 30 phyla by abundance are shown; all others are grouped as “Others”. The bar plot above summarizes the total number of detected propeller proteins per phylum. (H) Hierarchical cluster plot of all propeller proteins included in the family assignment analysis in panel (G), generated using ProteinClusterTools at bitscore cutoffs of 80, 120, 160, 200, 240, 280, and 320. Each circle represents a connected component (cluster) at a given bitscore threshold. Clusters at lower bitscore cutoffs (broader homology) form parent neighborhoods that progressively subdivide into smaller, more homogeneous clusters at higher cutoffs (higher homology). Each cluster is colored according to its dominant ECOD family. (I) Structural similarity of detected propellers to ECOD reference structures, shown as a hexbin plot of alignment length versus TM-score for the best Foldseek hit of each entry. Marginal distributions of both variables are shown. The vertical dashed line indicates a TM-score threshold of 0.5. Representative examples highlighting structural novelty are illustrated, including a 13-bladed propeller (AFDB entry: A0A2N2ZE83), a 14-bladed propeller (AFDB entry: A0A9D1M187), a propeller with a ‘lying’ meander conformation (AFDB entry: A0A9D7N7Y6). (J) UpSet plot comparing three propeller datasets derived from AFDB50: UniPropeller, TED propellers, and HMMER-based propeller annotations. Only regions with average pLDDT ≥ 75 were considered to ensure dataset quality. (K) Representative example (AFDB entry: A0A2A3F5H2) illustrating differences in residue coverage between UniPropeller and TED annotations, with corresponding structural regions shown. All structure models are colored by AlphaFold prediction confidence (pLDDT) throughout this study: dark blue (>90), sky blue (70–90), yellow (60–70), and orange (<60).

Utilizing a conservative segment extraction criterion (minimum score *S* = 0.8 and minimum length *L* = 20), we identified 799,263 (1.6%) proteins containing at least three predicted propeller blades. Among these detected proteins, the number of predicted blades most often corresponded to canonical propeller sizes, with proteins containing 7 predicted blades being most abundant, followed by proteins with 6, 5, 4, and 8 predicted blades, together accounting for 73.0% of all detected proteins (Figure 5B). In addition, 14.7% of entries contained three predicted blades, likely representing incomplete propeller domains. Proteins with 9 to 14 predicted blades were observed at lower but comparable frequencies, with entries containing 14 predicted blades being particularly enriched, presumably due to the tandem arrangement of two 7-bladed domains within a single chain.

Analysis of the within-protein blade sequence identity revealed a broad distribution centered at a mean of 28.6% (Figure 5C). Notably, 16.7% of these entries exhibited repeat identities below 20%, indicating a substantial fraction of highly divergent propellers. Conversely, 3.2% of entries displayed repeat identities above 60%, consistent with recently amplified propellers reported in previous studies (5). Evaluation of predicted structural confidence of these proteins showed a bimodal distribution of average pLDDT scores across blade regions (Figure 5D), with 76.7% of entries exceeding a mean pLDDT of 70. Functional annotation (Clusters of Orthologous Groups, i.e. COG) using eggNOG-mapper revealed that a considerable fraction of proteins (37.0%) lacked functional assignment (COG category ‘S’ and ‘-’; Figure 5E). Among annotated entries, the most common assigned categories were carbohydrate transport and metabolism (category ‘G’, 8.9%), intracellular trafficking, secretion, and vesicular transport (category ‘U’, 7.5%), signal transduction mechanisms (category ‘T’, 7.5%), posttranslational modification, protein turnover, chaperones (category ‘O’, 5.9%), and cell wall/membrane/envelope biogenesis (category ‘M’, 4.6%). From a taxonomic perspective, detected propeller-containing proteins were broadly distributed across the tree of life, with 54.4%, 42.0%, and 2.3% from Bacteria, Eukaryota, and Archaea, respectively, and the remaining 1.3% unclassified (Figure 5F). Within these three domains of life, the FCB group (13.5%) in Bacteria, as well as Metazoa (14.6%) and Fungi (9.7%) in Eukaryota, accounted for the largest numbers of propeller-containing proteins.

To improve dataset quality for downstream analyses, we applied a pLDDT threshold of 75, resulting in a high-confidence set, hereafter referred to as the UniPropeller dataset. For computational efficiency, UniPropeller sequences were further clustered at a maximum sequence identity of 50% (UniPropeller50). Assignment of UniPropeller50 sequences to ECOD propeller families using MMseqs2 revealed that the dataset encompasses a wide range of well-characterized β-propeller families (Figure 5G), including WD40 (23.7%), SGL (12.0%), DPPIV (9.4%), PQQ (8.1%), PSSI_BNR (6.5%), Kelch (5.8%), Lactonase (5.1%), RCC (4.1%), and VCBS (1.4%). Beyond these known families, clustering of the 393,255 UniPropeller50 sequences identified multiple clusters lacking significant similarity to any annotated ECOD propeller family (Figure 5H), highlighting substantial unannotated sequence diversity, as shown in SI Figure 17. For instance, one representative protein (AFDB entry: A0A9D7CAU2) adopts a canonical 7-bladed propeller topology, yet fails to produce any detectable hits in HHpred searches against ECOD70 (SI Figure 18).

We next examined the structural diversity of these confident (pLDDT ≥ 75) propeller-containing proteins by searching against ECOD propellers using Foldseek. 91.7% of entries exhibited clear structural similarity to known propellers, with TM-scores exceeding 0.5 and average alignment lengths of approximately 250 residues (Figure 5I). Notably, among the remaining domains with no clear structural matches to ECOD propeller families (TM-score < 0.5), we identified propellers with unusually large single-ring architectures (e.g., 13-bladed architecture A0A2N2ZE83; 14-bladed architecture A0A9D1M187) as well as domains exhibiting pronounced deviations in repeat-unit orientation (A0A9D7N7Y6, ‘lying conformation’). Despite these novel structural features, these propeller domains still yielded significant HHpred hits to annotated Pfam propeller families, matching Beta-prop_MIP2 (PF30506), DUF5046 (PF16465), and Conj_beta_prop_9 (PF27095) for A0A2N2ZE83, A0A9D1M187, and A0A9D7N7Y6, respectively (SI Figure 19-21). Existing eggNOG-mapper annotations provided little functional information about these proteins, with the exception of A0A9D1M187, which was annotated as a WG-containing protein and a putative RsiV-like protein. Despite these significant sequence-level matches, the representative proteins remained largely functionally uncharacterized, highlighting the functional darkness associated with structurally diverse β-propellers and motivating further investigation of these unusual repeat architectures.

Finally, we compared the UniPropeller50 dataset with propeller annotations generated using alternative approaches, including HMMER-based searches initiated from the same training propellers and the recent TED database, which assigns AFDB entries to CATH domains based on structural segmentation. Although this comparison is not strictly one-to-one, as TED annotations were generated using an earlier AFDB release while our scan was performed on the latest AFDB50 version, it nevertheless provides a qualitative assessment of relative coverage trends. After applying the same pLDDT ≥ 75 filter to all datasets, UniPropeller50 yielded broader coverage of propeller-containing proteins than either alternative approach (Figure 5J). Moreover, for proteins present in both UniPropeller50 and TED, the propellers identified by TREAD were often more extensive than their counterparts in TED (SI Figure 22). For example, in AFDB entry A0A2A3F5H2 (Figure 5K), TREAD predicted a complete 7-bladed propeller, whereas TED annotated only a partial 4-bladed domain, illustrating how sequence-based approaches can complement structure-based domain annotation pipelines.

### A cryptic connection links a novel barrel-like fold to propellers

Among the structurally atypical domains without similarity to ECOD propeller families, one class stood out by exhibiting a five-copy barrel-like fold composed of a repeated β1-α-β2-β3 unit, in which three β-strands form an anti-parallel β-meander that is interrupted by an α-helix at the sequence level (Figure 6A; representative AFDB entry: A0AAW5V244). HHpred searches against ECOD70 and PDB70 failed to detect clear homologous matches (SI Figure 23), except for a recently solved but still unclassified structure (PDB ID: 7TPU) featuring the same five-copy fold composed of β1-α-β2-β3 units. To assess whether this barrel-like architecture represents a larger family, we used the A0AAW5V244 structure as a query for Foldseek searches against AFDB50, retrieving a set of related proteins, visualized as a sequence similarity network in Figure 6B. This fold is distributed mostly in Bacteria and Eukarya, with a smaller number in Archaea, and, in addition to the five-copy form, also includes four-copy and six-copy architectures built from the same repeat unit, highlighting the internal structural diversity.

**Figure 6.**
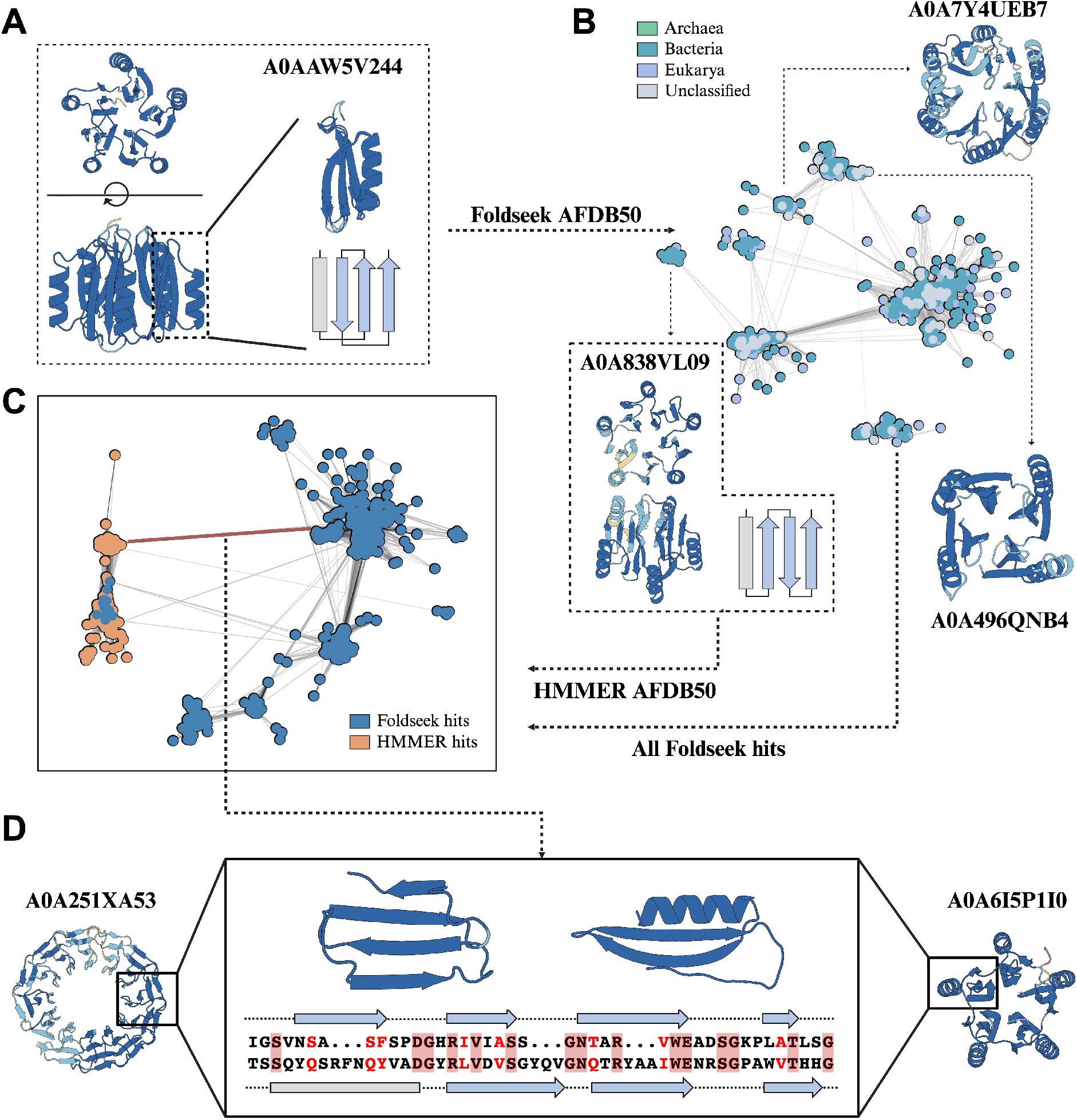
Cryptic connection between canonical propellers and a newly detected barrel-like fold. (A) Representative structure of the identified five-copy barrel-like fold (AFDB entry: A0AAW5V244). Secondary structure elements of a single repeat unit are illustrated in cartoon. (B) CLANS sequence similarity network generated from a Foldseek search against AFDB50 using the representative barrel-like domain as a seed (edge E-value threshold: 1×10^-3^). Nodes are colored according to taxonomic origin: Archaea (green), Bacteria (teal), and Eukarya (light purple). AFDB entries with unclassified taxonomy are shown in light grey. Clusters containing domains adopting the same repeat unit but organized as four-copy and six-copy architectures are illustrated using representative domains A0A496QNB4 and A0A7Y4UEB7, respectively. One cluster exhibits a decorated propeller topology, for which both the structure and the repeat unit topology of the representative domain A0A838VL09 are shown. (C) CLANS sequence similarity network generated from a HMMER search against AFDB50 using the propeller-like domain (A0A838VL09), combined with all proteins collected in (B) (edge E-value threshold: 1×10^-3^). Sequences retrieved by Foldseek and HMMER are shown in blue and orange, respectively. Directly detectable sequence similarity between canonical propellers and the barrel-like domain is observed, as indicated by the dark red edge. (D) Example pairwise alignment between a 12-bladed propeller (AFDB entry: A0A251XA53) and the five-copy barrel-like domain (AFDB entry: A0A6I5P1I0). The repeat-level alignment is extracted from the full-length alignment shown in Supplementary Figure 25, with identical residues shaded in dark pink and similar residues in red. Corresponding tertiary and secondary structures of the aligned repeat units are shown. All structure models are colored by AlphaFold prediction confidence (pLDDT): dark blue (>90), sky blue (70–90), yellow (60–70), and orange (<60).

Interestingly, among proteins retrieved with Foldseek (Figure 6B) we found a structural outlier (AFDB entry: A0A838VL09) whose repeat units adopt a propeller-blade-like topology (α-β1-β2-β3), resembling a canonical β-propeller blade in which the first β-strand is replaced by an α-helix. HHpred analysis of this domain (SI Figure 24) revealed high-confidence propeller matches, suggesting it may correspond to an intermediate that bridges canonical propellers and the barrel-like fold. To test this hypothesis, we combined sequences retrieved by HMMER searches initiated from this propeller-like domain with those previously identified by Foldseek for the barrel-like fold, and performed joint sequence clustering (Figure 6C). In the resulting map, the cluster containing proteins with the propeller-like topology (α-β1-β2-β3) was expanded with more examples, surprisingly including structures featuring the canonical propeller β1-β2-β3-β4 topology. This suggests that the α-β1-β2-β3 topology does not form a separate intermediate subfamily, but possibly can be interchanged to the canonical propeller β1-β2-β3-β4 topology with a limited number of mutations. Multiple significant pairwise alignments were detected between the mixed propeller/propeller-like cluster and sequences corresponding to the β1-α-β2-β3 topology. One representative example shows significant sequence similarity between a 12-bladed propeller and the barrel-like fold (SI Figure 25, E-value = 1×10^−13^, 30% sequence identity, and 39% similarity), with a unit-level alignment shown in Figure 6D. The unit alignment reveals a coherent correspondence in terms of secondary structure: the first β-strand of the 12-bladed propeller unit aligns to the α-helix of the barrel-like fold; the second and third propeller β-strands align to two consecutive β-strands in the other fold; and the last propeller β-strand aligns to the first β-strand of the next repeat in the barrel-like fold. Conserved residues occur at comparable positions in both units, including a GN motif at the tip of β3. Notably, the Velcro-like closure, a hallmark of most propellers, is also present in this five-copy repetitive domain. Despite the significant sequence similarity detected in such pairs, the functional role of this five-copy barrel-like domain still remains unclear. In the experimentally solved polyglycine hydrolase from *Fusarium vanettenii* (PDB ID: 7TPU), it forms an N-terminal domain adjacent to the catalytic β-lactamase-like serine protease domain (60). A subsequent mutagenesis and docking study argued against a direct role of this domain in substrate recognition or catalysis (61). Together with our observation that this fold occurs either as standalone domains or fused to diverse functional domains, frequently proteases or kinases, these findings suggest that this novel barrel-like fold may instead serve as a modular scaffold involved in protein–protein interactions, regulation, or recruitment and stabilization of associated domains.

Together, these observations identify a repeat-unit–level connection linking a previously unclassified barrel-like fold to canonical propellers. While this analysis does not completely establish an evolutionary relationship between the barrel-like fold and canonical propellers, it demonstrates that repeat-unit similarity can persist across proteins with markedly different global architectures. More broadly, it illustrates how large-scale, unit-resolved repeat annotation driven by embedding-based methods can expose cryptic connections that remain difficult to detect using traditional analyses.

### β-propellers show lineage-specific expansion and functional deployment across the tree of life

Having established a comprehensive and high-confidence census of β-propeller architectures at the protein level, we next asked how these domains are distributed across the tree of life, and whether their functional composition differs between lineages. To address this, we applied the developed workflow to a curated set of 4,618 representative proteomes spanning major evolutionary lineages (12). Across proteomes, the fraction of proteins containing at least three predicted propeller blades showed a moderate correlation with proteome size (SI Figure 26; Spearman’s R=0.330; p < 0.001), but also exhibited pronounced lineage-specific patterns. In particular, eukaryotic genomes encode substantially higher proportions of propeller-containing proteins than either bacterial or archaeal genomes (Figure 7A-B).

**Figure 7.**
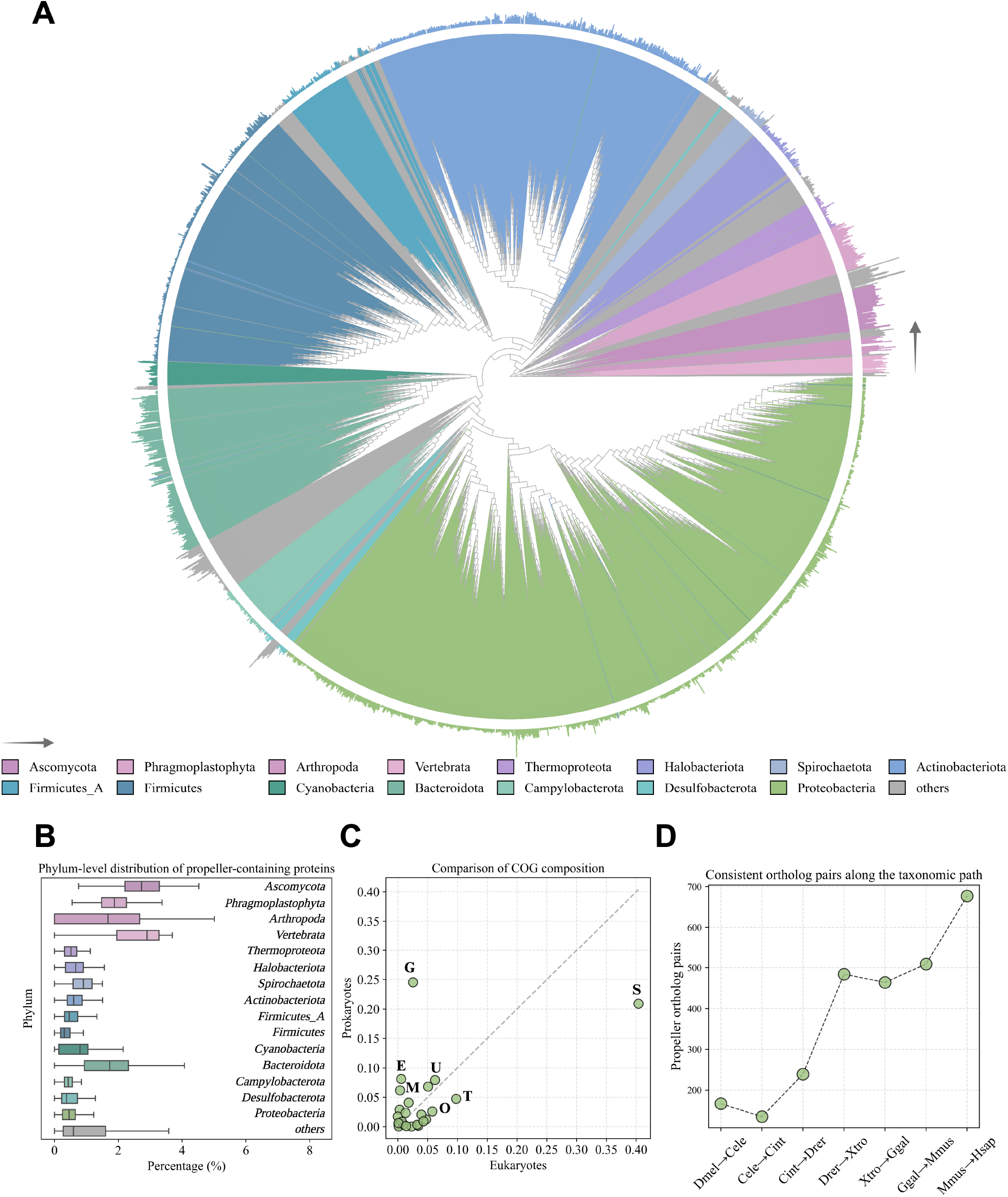
Propeller model scan on representative proteomes spanning the tree of life. (A) Phylogenetic tree of 4,618 representative species, retrieved from Cho et al., with branches colored by phylum. For each species, the proportion of propeller-containing proteins (defined as proteins containing at least three propeller blades) relative to proteome size is shown as an outer-ring bar plot. Phylum colors follow the legend, arranged in clockwise order starting from *Ascomycota*, corresponding sequentially to the left-to-right order of phyla in the legend. (B) Phylum-level distribution of propeller-containing proteins across representative proteomes, shown using the same color scheme as in panel (A). (C) Comparison of COG functional composition between propeller-containing proteins detected in Bacteroidota and those from other bacterial phyla. COG categories displaying pronounced compositional differences are labeled with the COG letter. G: Carbohydrate transport and metabolism; T: Signal transduction mechanisms; S/-: Function unknown; E: Amino acid transport and metabolism; U: Intracellular trafficking, secretion, and vesicular transport; M: Cell wall/membrane/envelope biogenesis; O: Posttranslational modification, protein turnover, chaperones. (D) Consistent ortholog pairs containing propeller proteins along the taxonomic path: *Drosophila melanogaster*, *Caenorhabditis elegans*, *Ciona intestinalis*, *Danio rerio*, *Xenopus tropicalis*, *Gallus gallus*, *Mus musculus*, and *Homo sapiens*.

To investigate whether these differences in abundance are accompanied by functional specialization, we annotated propeller-containing proteins and analyzed their COG composition (SI Table 5). In eukaryotes, a large fraction of propeller-containing proteins was assigned to poorly characterized / unknown categories (COG R/S/-; 40.4%), compared with 20.9% in prokaryotes (Figure 7C). Beyond this difference, several well-defined functional categories exhibited pronounced lineage-specific biases. Prokaryotic propeller domains were overrepresented in carbohydrate transport and metabolism (COG G; 24.5% versus 2.5% in eukaryotes), as well as in cell wall, membrane, and envelope biogenesis (COG M; 6.2% versus 0.4%) and amino acid transport and metabolism (COG E; 8.1% versus 0.6%). In contrast, eukaryotic propeller-containing proteins showed higher contributions to posttranslational modification, protein turnover, and chaperone-related functions (COG O; 5.8% versus 2.6% in prokaryotes), as well as to signal transduction mechanisms (COG T; 9.8% versus 4.7%).

Despite the overall lower abundance of propeller-containing proteins in prokaryotic proteomes, one bacterial lineage, Bacteroidota, emerged as a notable outlier. In this phylum, nearly 2% of proteins per proteome contained at least three propeller blades, approaching the fraction observed in eukaryotic proteomes (Figure 7B). While both Bacteroidota and other bacterial phyla predominantly deploy propeller domains in carbohydrate-related processes (COG G; 24.0% and 25.4%, respectively; SI Figure 27), Bacteroidota showed a markedly higher representation of propeller domains in signal transduction functions (COG T; 10.6% compared with 3.0% in other bacteria, SI Figure 27). This bias toward signal transduction-related functions is consistent with the ecological roles of Bacteroidota as highly responsive environmental generalists, which rely extensively on regulatory and sensing systems to adapt to fluctuating nutrient landscapes in animal-associated and other nutrient-variable habitats, such as the gut (62). Among prokaryotic phyla with many detected propellers, Cyanobacteria displayed a distinct functional bias of propeller-containing proteins. Compared with other prokaryotes, they showed a markedly reduced representation in carbohydrate-related processes (COG G; 9.0% versus 25.5%), consistent with their photoautotrophic lifestyle. Conversely, propeller domains were enriched in transcription and DNA replication/repair functions (COG K and L; 9.0% and 8.3% versus 1.9% and 3.9%), suggesting a preferential association with regulatory and genome maintenance processes, potentially linked to light-responsive physiology.

Finally, to examine propeller conservation along increasing organismal complexity, we analyzed orthologous protein pairs across a representative series of metazoan species, including *Drosophila melanogaster*, *Caenorhabditis elegans*, *Ciona intestinalis*, *Danio rerio*, *Xenopus tropicalis*, *Gallus gallus*, *Mus musculus*, and *Homo sapiens*. For each pair of consecutive species, orthologs were classified as potentially conserved propeller pairs if both proteins contained at least three detected propeller blades. The number of such ortholog pairs increased progressively along this taxonomic path (Figure 7D), consistent with increasing prevalence of propeller-containing ortholog pairs along this metazoan lineage.

Together, these analyses place the large-scale propeller landscape uncovered by TREAD into an evolutionary and functional context, revealing pronounced lineage-specific patterns in both the abundance and deployment of β-propeller domains across the tree of life.

## Discussion

In this study, we demonstrate that protein repeat detection can be reformulated as a supervised residue-level annotation problem using TREAD. Instead of encoding repeat family information into explicit probabilistic profiles, TREAD captures repeat-specific features implicitly through discriminative learning on protein embeddings. Across two complementary benchmark scenarios, this framework consistently matched or exceeded the performance of the profile-based HMMER baseline, while offering greater flexibility in task design. We chose HMMER as the representative profile-based method because it uses prior knowledge (as does TREAD), provides a widely adopted, general-purpose engine underlying major annotation resources such as Pfam and SMART (21, 63), and offers a highly reproducible and scalable pipeline for building and scanning profile databases. We did not include HHpred in the main benchmarks, as its reliance on query-specific profile construction would substantially increase the computational cost of evaluating the full RepeatsDB and Pfam datasets. Alternative profile-based repeat detection tools, such as REP and TPRpred (22–24), are also available, but their method-specific design choices make them less suitable as family-agnostic baselines for systematic benchmarking across diverse repeat types. Also, we did not include *de novo* approaches in the main benchmarks, as they represent a different class of repeat detection methods; however, we performed an additional comparison with such methods (SI Figure 6), which supported the advantage of prior-knowledge-driven approaches when the target repeat type is predefined.

At its core, TREAD treats repeat detection as a residue-level discrimination between repeat and non-repeat, similar to other residue-wise prediction tasks such as ligand-binding site or post-translational modification prediction (64, 65). The effectiveness of this formulation likely reflects two sources of contextual information. First, residue embeddings derived from pretrained pLMs encode contextual information shaped by evolutionary constraints, which explains why a simple linear layer achieved reasonable performance in ablation experiments (SI Table 2). Second, the task-specific model architecture further integrates contextual information through convolutional and recurrent components, enabling the model to refine local residue-level patterns using broader motif context. This additional contextual modeling appears particularly beneficial for alignment-poor datasets or highly diverged repeat families. The superior performance of TREAD over HMMER cannot be attributed solely to these two features, as the methods also differ fundamentally in their learning paradigms. However, the fact that the unsupervised *de novo* method pLM-Repeat, which also utilizes ProtT5 embeddings, performed worse than TREAD (SI Figure 6) suggests that the superior performance of TREAD may partly reflect the benefits of task-specific supervised learning.

Another advantage of the TREAD framework is its flexibility with respect to both training data and task formulation. Profile-based methods require well-curated alignments for each targeted repeat family, whereas TREAD can be trained on any dataset that provides residue-level annotations, regardless of whether these annotations originate from structural databases, expert curation, or automated pipelines. This decoupling of repeat annotation from explicit alignment construction allows the framework to operate in settings where homology-based profile construction is difficult or impractical. In addition, the framework naturally supports multi-task or multi-class repeat annotation through modification of the prediction heads as illustrated in the RepeatsDB evaluation, where multiple repeat protein folds were considered simultaneously.

Importantly, this flexibility also means that the biological scope of a TREAD model is determined by the definition and diversity of its training labels. Broadly defined labels can be used to learn repeat features shared across heterogeneous sequence, structural, and functional contexts, whereas more narrowly defined labels yield correspondingly more specific annotation models. This distinction is illustrated by the RepeatsDB model, which learned six broad repeat-fold classes, in contrast to the Pfam models trained separately for individual repeat families. Similarly, the propeller blade model was deliberately trained across diverse β-propeller families to capture blade-level features shared broadly across the fold. Consequently, a shared TREAD annotation should be interpreted at the biological level represented by its training labels and does not by itself imply similarity in full-length protein architecture, function, or evolutionary history.

Although TREAD is presented here for protein repeat annotation, the framework is readily extensible to other motif-centric residue-level prediction tasks. A related idea has already been explored for Rossmann-like nucleotide-binding motifs (66), whereas other potential applications include motifs such as Walker-A P-loop motifs, as well as structurally defined motifs (e.g., coiled-coils or helix-turn-helix motifs) given appropriate residue-level annotations. Moreover, TREAD is agnostic to the choice of input representation and can readily incorporate diverse pretrained residue-level embeddings, ranging from pLM embeddings to structure-based and emerging multimodal representations (67, 68). Together, this flexibility makes TREAD a general interface for leveraging transfer learning in supervised residue-level motif annotation tasks.

Beyond benchmarking, the β-propeller analysis illustrates how an embedding-based framework can provide analytical perspectives that are difficult to obtain using conventional profile-centric approaches alone. Applying the propeller-specific TREAD model to the AFDB50 produced a comprehensive census of the propeller landscape, yielding the UniPropeller dataset. A notable outcome was the identification of a previously uncharacterized five-fold barrel-like repeat fold that is not readily captured by standard homology detection methods. At the repeat-unit level, statistically significant similarity is observed between this fold and canonical propeller blades, consistent with a possible unit-level evolutionary relationship. These observations motivate further investigation of repeat fold relationships despite differences in repeat-unit topology. Moreover, extending the propeller TREAD model across representative proteomes placed these findings in an evolutionary context. The pronounced enrichment of propeller-containing proteins in eukaryotic proteomes, and their preferential association with regulatory and post-translational functions, contrasts with the more specialized deployment observed in prokaryotes, particularly in carbohydrate-related and envelope-associated processes. The expansion observed in Bacteroidota shows how repeat architectures can be differentially amplified and repurposed in response to ecological and functional pressures.

Despite its strengths, the proposed approach has clear limitations. By design, it decouples repeat annotation from alignment construction and therefore does not directly provide repeat-to-repeat alignments, which can provide insights into conservation and variation within repeat units. Repeat alignments can currently only be obtained as a *post hoc* step by extracting predicted segments and aligning them separately. Addressing this limitation would require extending the framework to jointly model segmentation and unit correspondence, for example by incorporating differentiable alignment modules tailored to repeat unit matching (34, 69, 70).

Once a TREAD model has been trained or a repeat profile constructed, the cost of scanning individual query sequences is broadly comparable between the two approaches. Profile-based methods incur substantial upfront costs associated with homolog retrieval to construct alignments, whereas TREAD shifts this cost to model training and embedding generation. In the RepeatsDB benchmark, for example, training the model, including embedding generation, required approximately one GPU hour. In contrast, building MSAs for all repeat proteins in this benchmark consumed on the order of ∼200 CPU hours. At very large scales, the dominant bottleneck for TREAD shifts to embedding computation and batched model inference. For example, the AFDB50 scan required on the order of ∼1000 GPU hours. In contrast, HMMER remains highly optimized for certain large-scale search tasks. Thus, the choice between the two approaches depends on the availability of curated homolog data, the target scale of analysis, and the available computational resources.

In summary, this study shows that protein repeat annotation can be effectively reformulated as a supervised, residue-level learning problem operating directly on protein embeddings. By replacing explicit profile construction with discriminative models trained on annotated data, TREAD provides a flexible alternative to classical profile-based approaches while retaining sensitivity to weak evolutionary signals. Rather than competing with alignment-driven methods, this framework complements them by enabling scalable and fine-grained repeat annotation in settings where curated alignments are limited or impractical. More broadly, our results suggest that repeat proteins can be studied through a general motif-centric approach, opening new avenues for their systematic analysis beyond traditional alignment-based paradigms.

### Data availability

Source code of TREAD and a user-friendly command-line interface are available on the GitHub repository https://github.com/KYQiu21/TREAD/tree/main. The Colab Notebook for running models obtained in this study is available at https://colab.research.google.com/drive/1gbtb5BtevWE9vChJrYgiNW2mQSW_kN8j. The benchmark dataset, together with the trained model weight and the generated dataset are available on the Zenodo repository https://zenodo.org/records/20593717.

## Supplementary material description

S1_File: the supplementary materials including 27 supplementary figures and 5 supplementary tables. S2_File: the table including all detected repeat proteins by TREAD from the AFDB Swiss-Prot scan. S3_File: the table including all UniPropeller50 proteins and analyzed results from the AFDB50 scan. S4_File: the HTML file of the hierarchical cluster plot of UniPropeller50 proteins.

## Acknowledgements

Computations were performed on the MPI-BIO cluster and the HPC system Raven at the Max Planck Computing and Data Facility. K.Q. would like to thank the IMPRS From Molecules to Organisms PhD program. K.Q. thanks Tim Z. Xiao and Natalie Ditz for insightful discussions. This work was supported by institutional funds of the Max Planck Society. S.D.-H. was additionally supported by the National Science Centre (grant 2020/37/B/NZ2/03268).

