## Supplemental Materials for "Beyond profiles: supervised repeat annotation using protein embeddings"

### Supplementary Figures

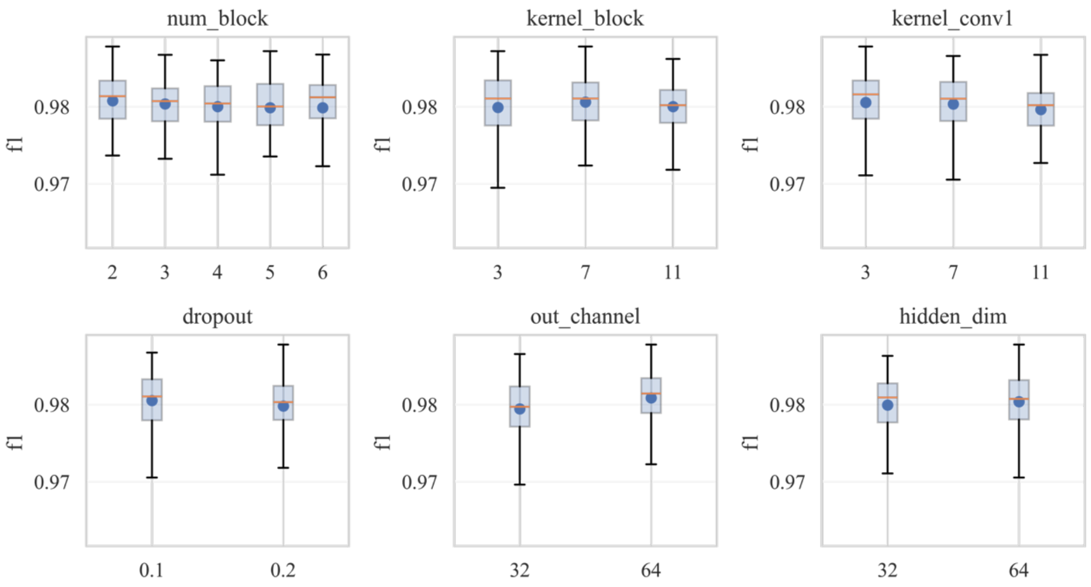


**Supplementary Figure 1:** Grid search of hyperparameters for TREAD model training on the RepeatsDB first-fold dataset. All models obtained from the grid search were evaluated on the same validation set using residue-level F1-score (see Materials and Methods for details). Six hyperparameters were explored, one per subplot: *num_block*, *kernel_block*, *kernel_conv1*, *dropout*, *out_channel*, and *hidden_dim*. Each box plot summarizes the distribution of F1-scores across all tested settings for a given hyperparameter, with the mean and median indicated by the orange line and blue dot, respectively.

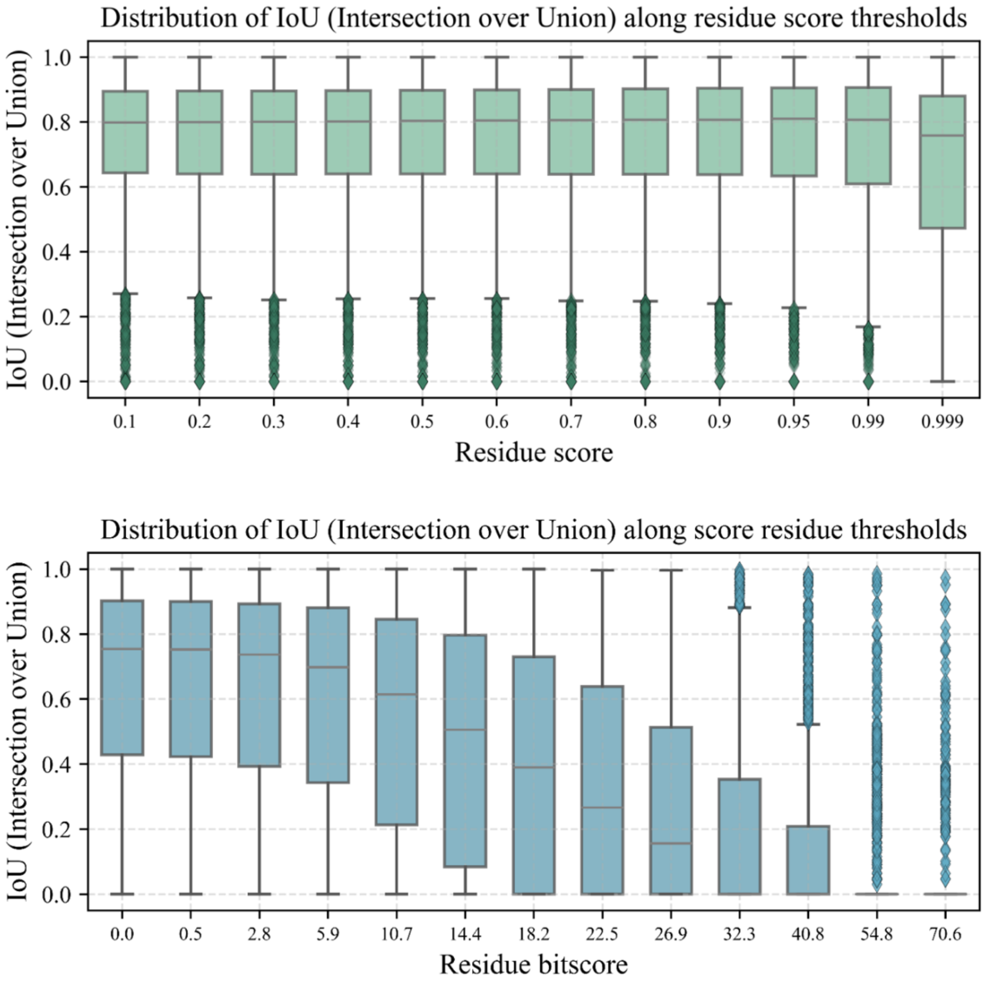


**Supplementary Figure 2:** Distribution of Intersection over Union (IoU) scores for TREAD (top) and HMMER (bottom) across residue score thresholds at the minimal length threshold ($L$) of 20. For each residue score threshold, IoU scores were computed between predicted and reference repeat segments and summarized as box plots. Boxes indicate the interquartile range (25th–75th percentiles), with whiskers extending to 1.5× the interquartile range. The median is shown as a solid black line, and outliers are indicated by diamond markers.

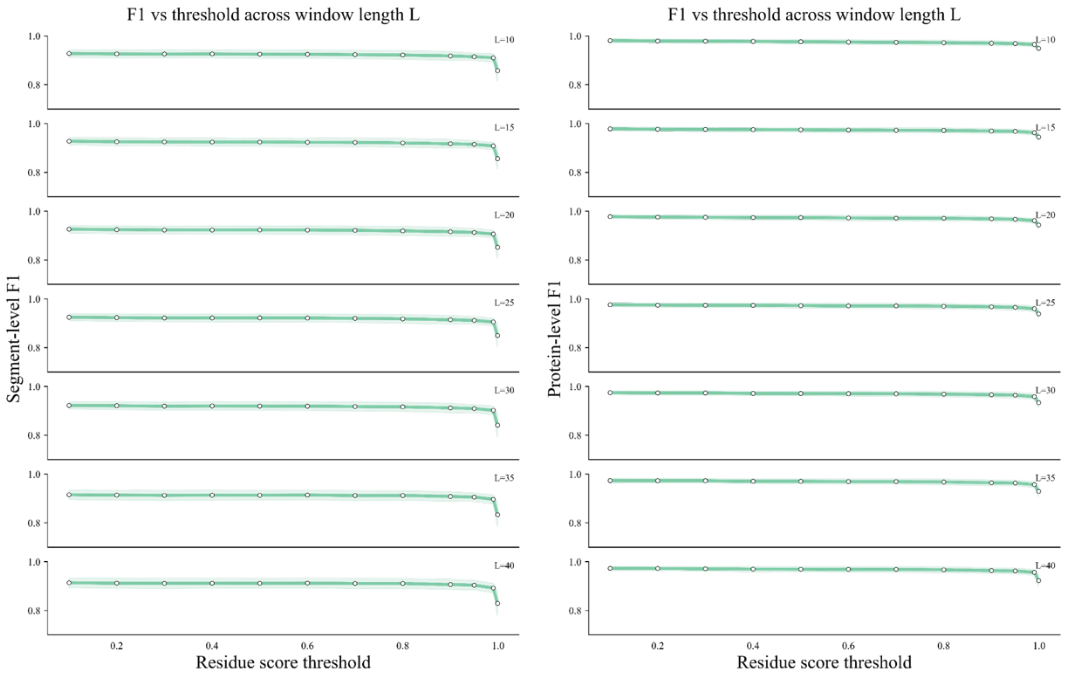


**Supplementary Figure 3:** Segment-level (left) and protein-level (right) F1-score of the TREAD model on RepeatsDB across residue score thresholds and minimum segment lengths. Performance was evaluated on independent test sets from five cross-validation folds. For each minimum segment length, a series of residue score thresholds were tested, following the procedure described in Figure 1C–D. Solid white points indicate the mean F1-score for each parameter combination, while shaded regions represent ±1 standard deviation across the five folds.

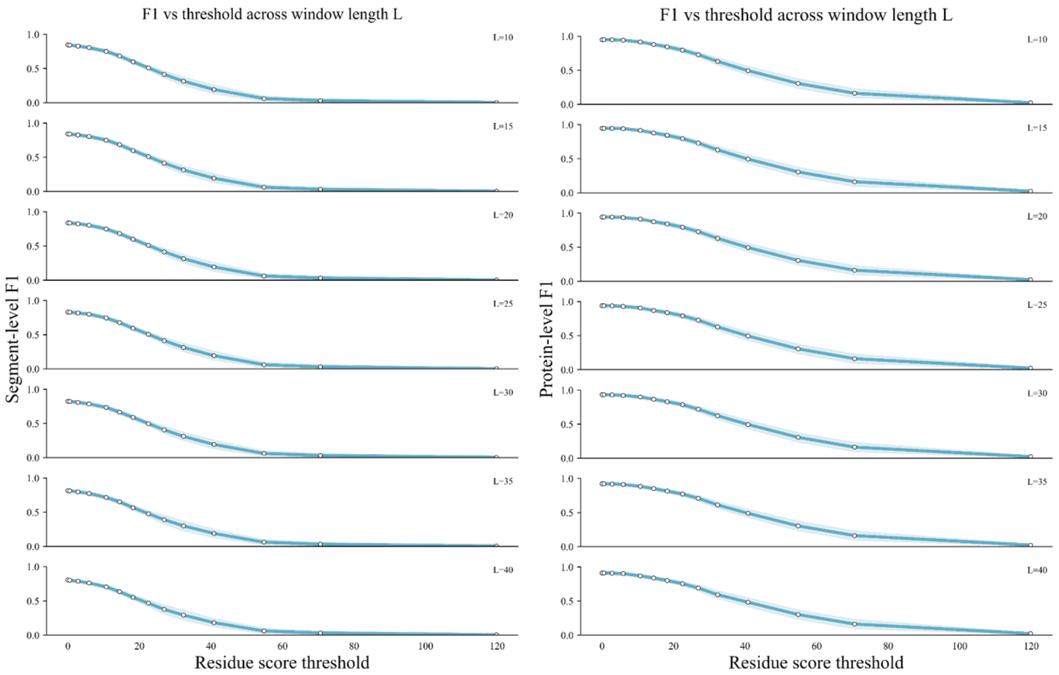


**Supplementary Figure 4:** Segment-level (left) and protein-level (right) F1-score of HMMER on RepeatsDB across residue score thresholds and minimum segment lengths. Performance was evaluated on independent test sets from five cross-validation folds. For each minimum segment length, a series of residue score thresholds were tested, following the procedure described in Figure 1C–D. Solid white points indicate the mean F1-score for each parameter combination, while shaded regions represent ±1 standard deviation across the five folds.

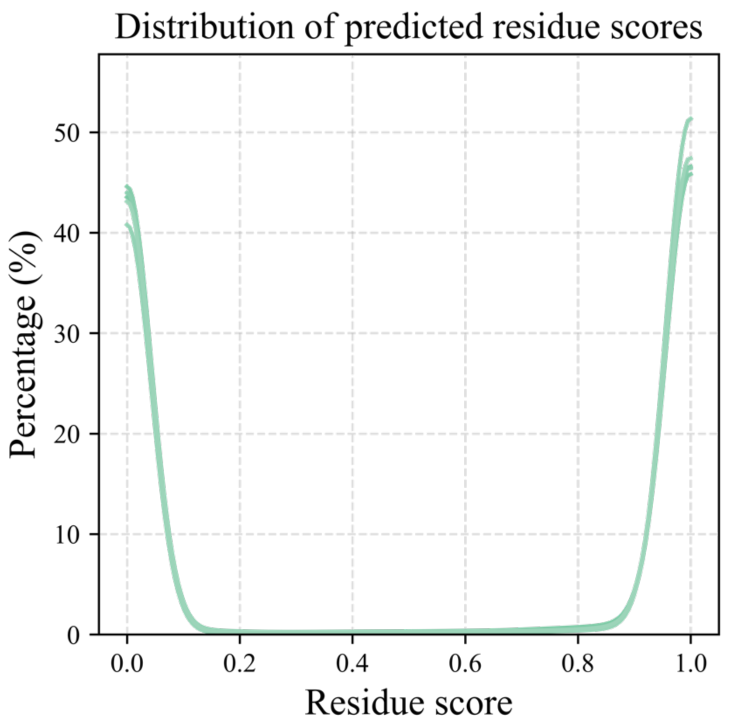


**Supplementary Figure 5:** Distribution of predicted residue scores for the positive examples in the test datasets of the RepeatsDB benchmark. Each curve represents one cross-validation fold. Residue scores were obtained from the prediction head corresponding to the ground-truth repeat fold.

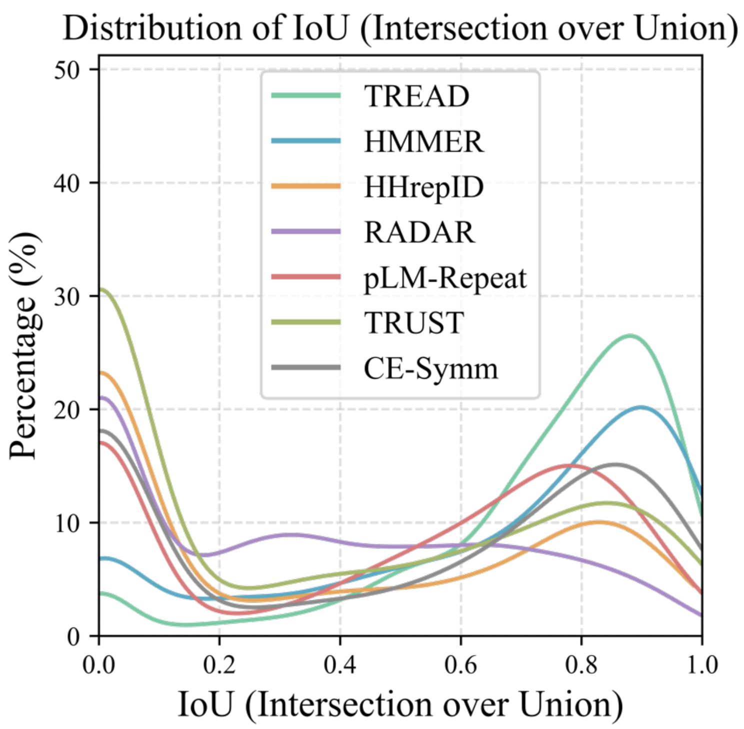


**Supplementary Figure 6:** Distribution of Intersection over Union (IoU) scores for repeat detection and annotation methods. The following thresholds were used: TREAD residue score threshold = 0.1; HMMER bit score threshold = 0.0; HHrepID P-value threshold = 0.001; pLM-Repeat self-alignment score threshold = 0.3; RADAR repeat family score threshold = 100; TRUST repeat family P-value threshold = 0.01; and CE-symm self-alignment score threshold = 0.4. Each method is represented by a single curve. For the two prior-knowledge-based methods, TREAD and HMMER, the curve was obtained by summing the five fold-specific curves from the independent cross-validation folds (Figure 1C). For the *de novo* repeat detection methods, the curve represents the distribution calculated from the RepeatsDB entries pooled across the five test datasets used for TREAD and HMMER. Only IoU scores from repeat-containing sequences are included.

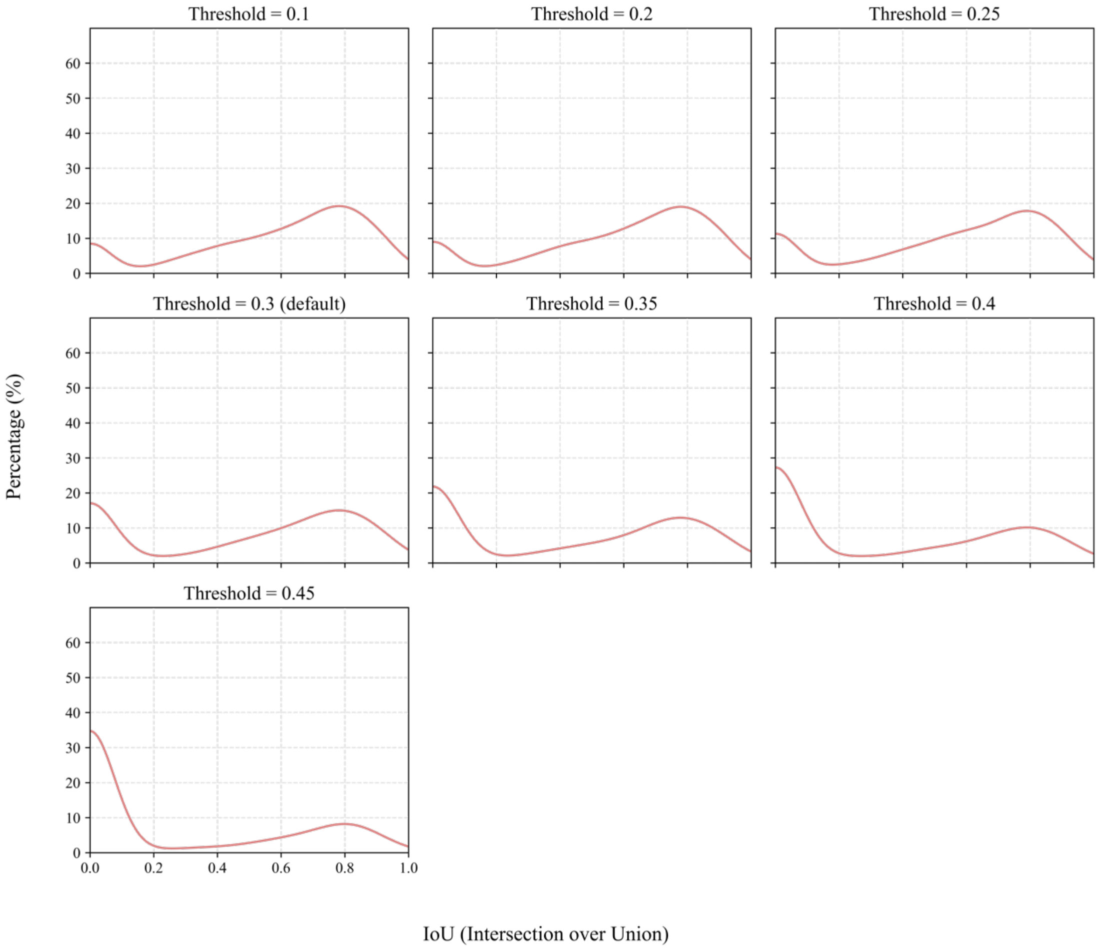


**Supplementary Figure 7:** Distribution of Intersection over Union (IoU) scores for pLM-Repeat across a series of self-alignment score thresholds: 0.1, 0.2, 0.25, 0.3, 0.35, 0.4, and 0.45. Higher thresholds correspond to more stringent detection criteria.

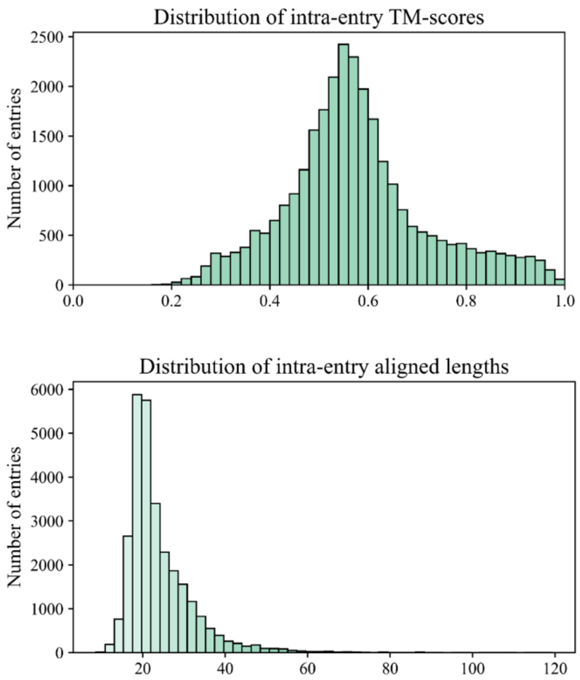


**Supplementary Figure 8:** Distribution of intra-entry TM-scores (upper) and aligned lengths (lower) for additional TREAD predictions in the AFDB Swiss-Prot dataset that were missed by STRPSearch. For each entry, the predicted segments were extracted from the corresponding AlphaFold Model and compared pairwise using TM-align.

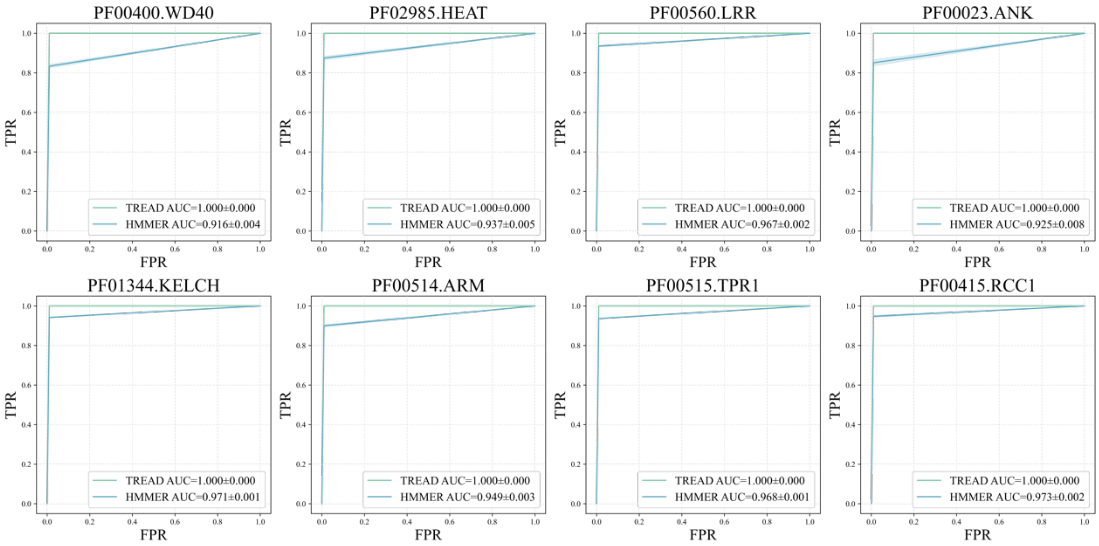


**Supplementary Figure 9:** Residue-level ROC curves for TREAD and HMMER across the eight Pfam families, evaluated on test datasets derived from Pfam full alignments. Plotting conventions are identical to Figure 1B.

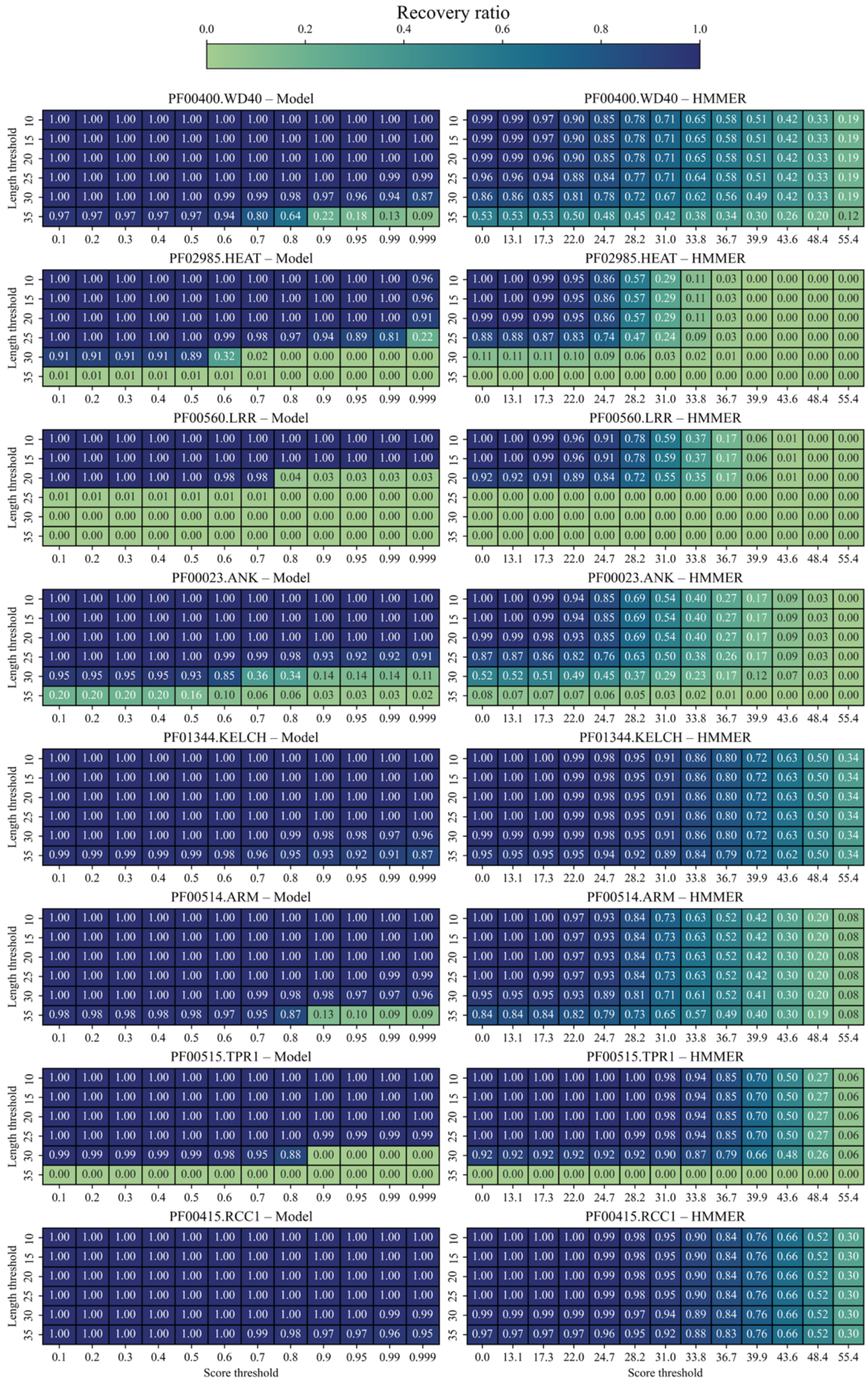


**Supplementary Figure 10:** Heatmap of segment-level F1-score for TREAD (left) and HMMER (right) across residue score thresholds and minimum segment lengths. Performance was evaluated on test sets extracted from Pfam full alignments for each Pfam repeat family. Each heatmap summarizes model performance across all parameter combinations for a given repeat type. A unified color scale is used across all panels, with higher F1-scores indicated by darker blue.

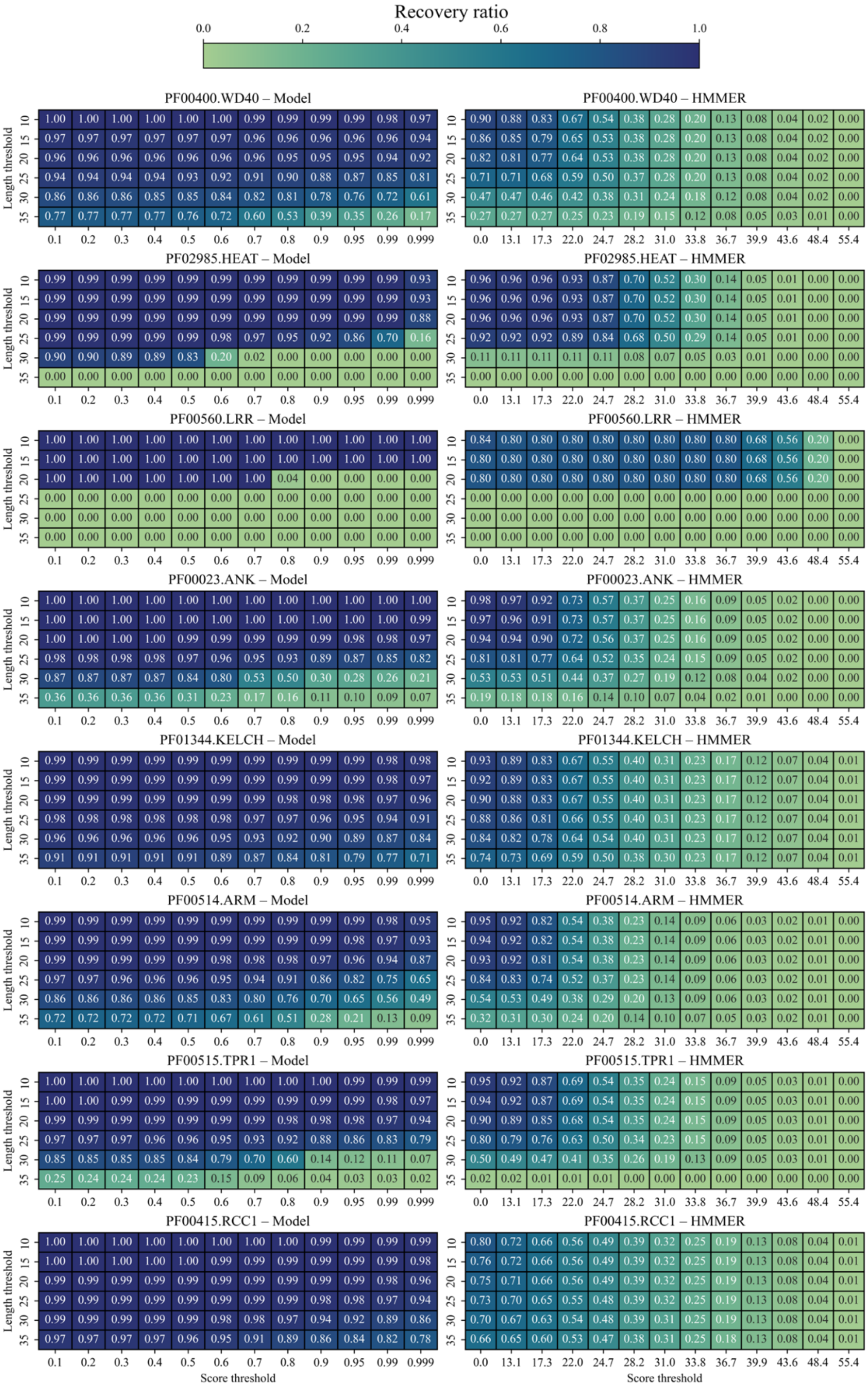


**Supplementary Figure 11:** Heatmap of segment-level Recall for TREAD (left) and HMMER (right) across residue score thresholds and minimum segment lengths. Performance was evaluated on PSI-BLAST–generated datasets for each Pfam repeat family. Each heatmap summarizes model performance across all parameter combinations for a given repeat type. A unified color scale is used across all panels, with higher Recall indicated by darker blue.

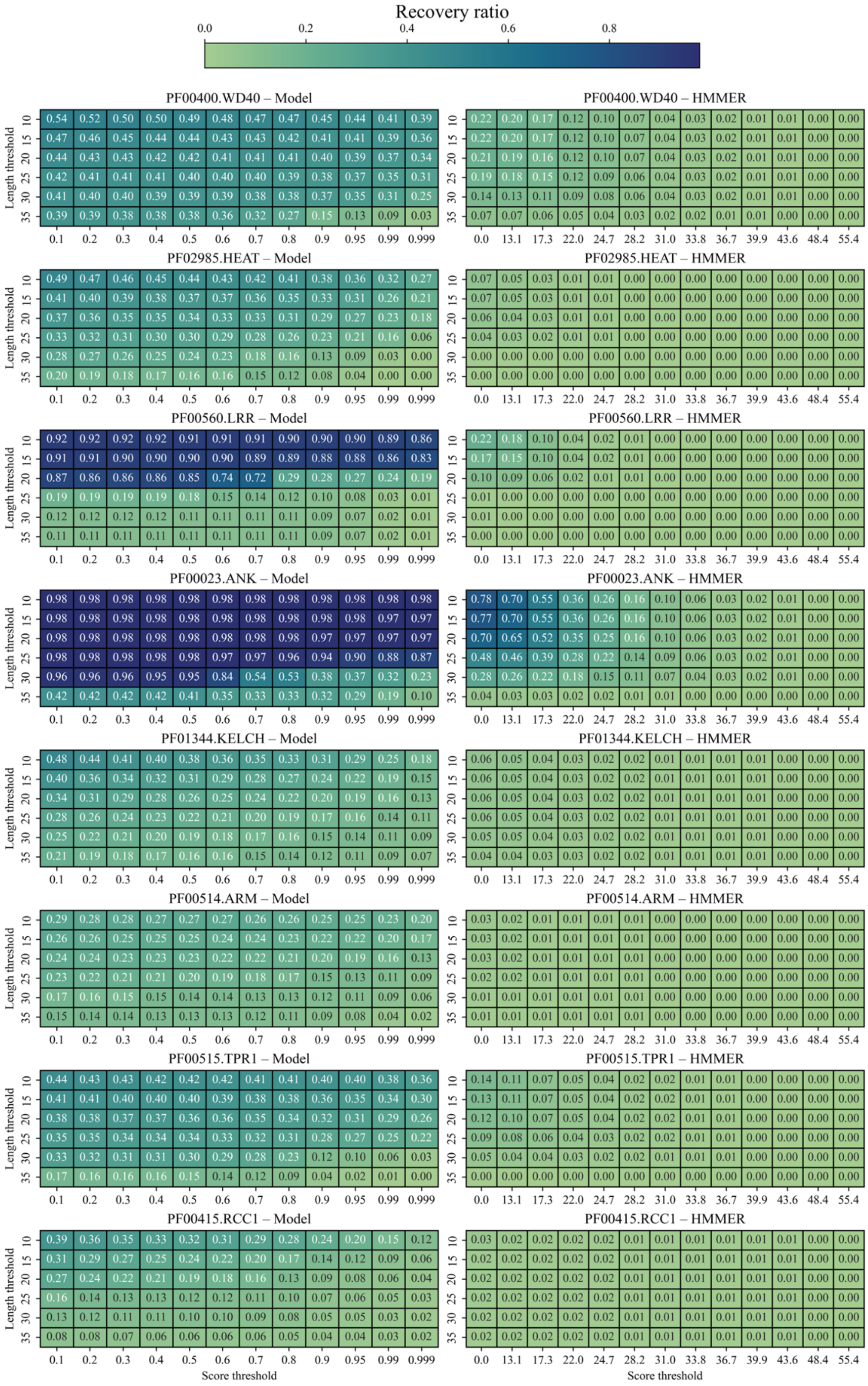


**Supplementary Figure 12:** Heatmap of F1-score under the relaxed presence/absence criterion for TREAD (left) and HMMER (right) across residue score thresholds and minimum segment lengths. Performance was evaluated on Pfam other-family datasets for each Pfam repeat family. Each heatmap summarizes model performance across all parameter combinations for a given repeat type. A unified color scale is used across all panels, with higher F1-scores indicated by darker blue.

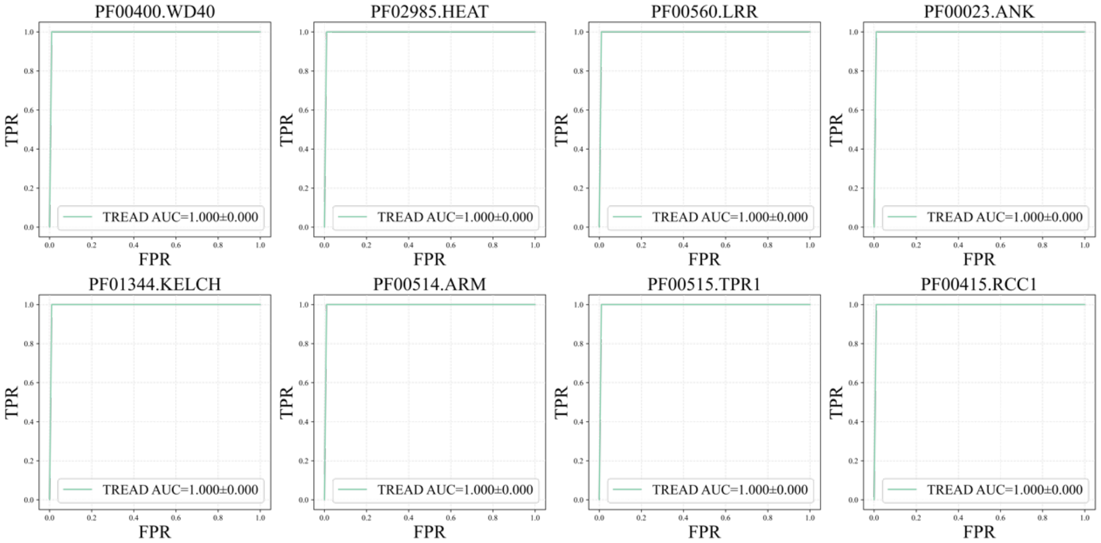


**Supplementary Figure 13:** Residue-level ROC curves for TREAD (linear) across the eight Pfam families, evaluated on the test datasets derived from Pfam full alignments. Plotting conventions are identical to Fig. 3B.

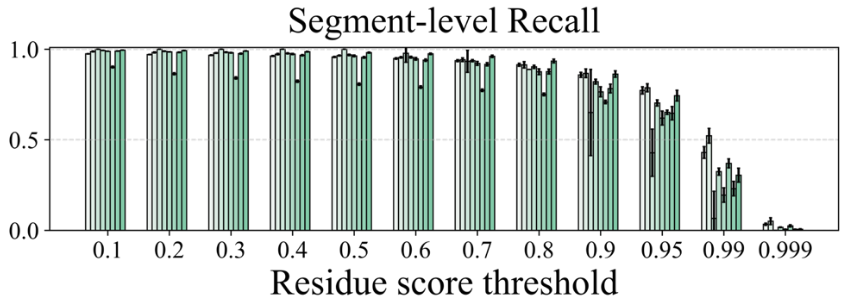


**Supplementary Figure 14:** Segment-level Recall across residue score thresholds for TREAD (linear) at the minimum repeat length of 20, evaluated on the PSI-BLAST-generated datasets.

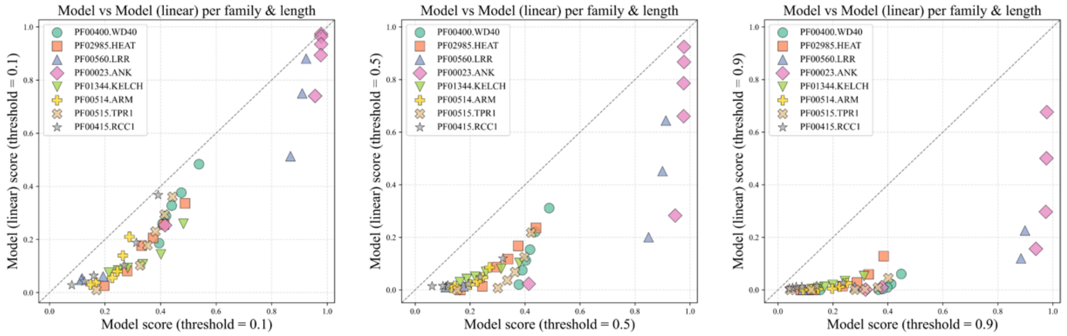


**Supplementary Figure 15:** Protein-level recall comparison between TREAD with full architecture and a linear-only variant. Performance was evaluated on datasets curated from Pfam families belonging to the same Pfam clan for each repeat type. Results are shown at residue score thresholds of 0.1 (left), 0.5 (middle), and 0.9 (right). For each repeat family, multiple minimum segment length thresholds (L = 10, 15, 20, 25, 30, and 35) were tested and plotted as individual points, with repeat families distinguished by unique marker shapes and colors.

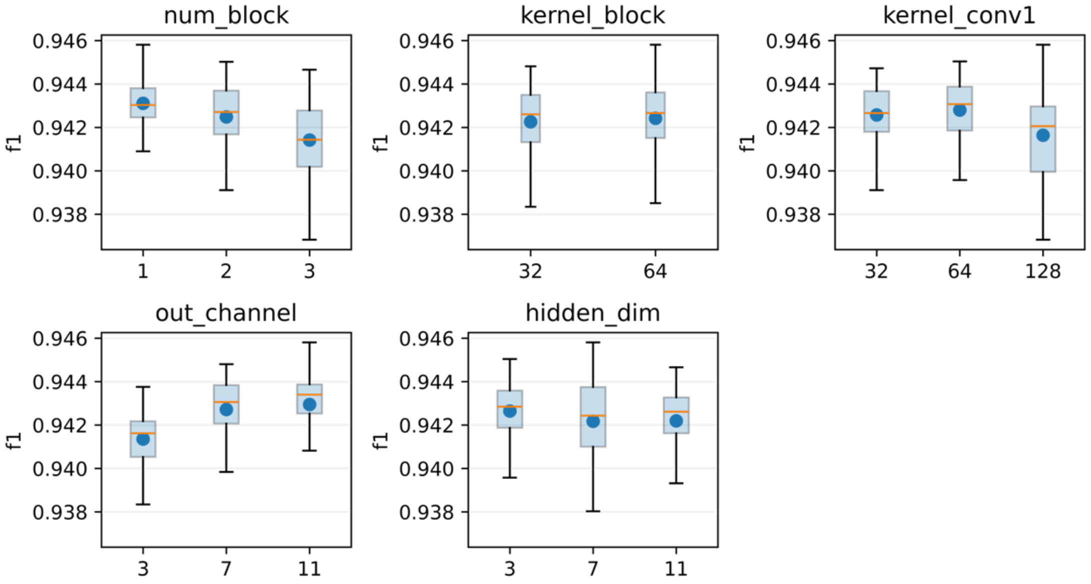


**Supplementary Figure 16:** Grid search of hyperparameters for TREAD model training on the propeller first-fold dataset. All models obtained from the grid search were evaluated on the same validation set using residue-level F1-score (see Materials and Methods for details). Five hyperparameters were explored, one per subplot: *num_block*, *kernel_block*, *kernel_conv1*, *out_channel*, and *hidden_dim*. Each box plot summarizes the distribution of F1-scores across all tested settings for a given hyperparameter, with the mean and median indicated by the orange line and blue dot, respectively.

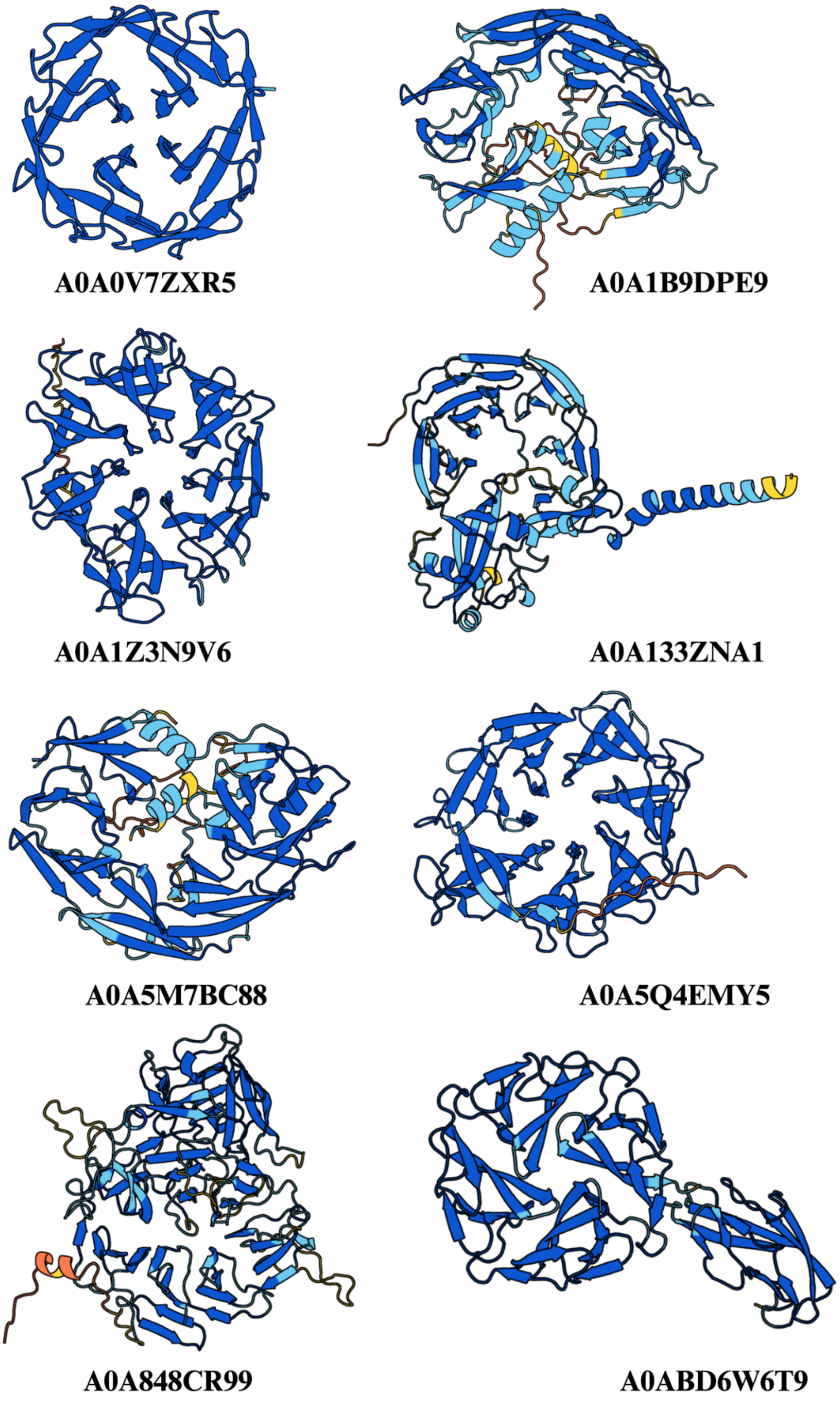


**Supplementary Figure 17:** Representative examples of detected propeller domains annotated as 'undefined cluster' without significant MMSeqs2 hit against ECOD propeller families.

```
Query         Q_5657867_1
Match_columns 377
No_of_seqs    231 out of 1762
Neff          9.92915
Searched_HMMs 71734
Date          Tue Feb 10 11:24:24 2026
Command       hhsearch -cpu 8 -i ../results/full.a3m -d /cluster/toolkit/production/databases/hh-
suite/ECOD/ECOD_F70_20230309 -o ../results/5657867_1.hhr -oa3m ../results/5657867_1.a3m -p 20 -Z 250
-loc -z 1 -b 1 -B 250 -ssm 2 -sc 1 -seq 1 -dbstrlen 10000 -norealign -maxres 32000 -contxt /cluster/
toolkit/production/bioprogs/tools/hh-suite-build-new/data/context_data.crf

 No Hit                             Prob E-value P-value  Score    SS Cols Query HMM  Template HMM
  1 ECOD_000005607_e2jnaA1 | 872.4   4.4 7.3E+02    0.01   18.7   1.2   19    1-19      1-19  (96)

No 1
>ECOD_000005607_e2jnaA1 | 872.4.1.1 | 2JNA A:1-96 | A: a+b two layers, X: Dodecin subunit-like, H:
NO_H_NAME, T: YdgH-like, F: DUF1471 | Protein: Putative secreted protein
Probab=4.37  E-value=7.3e+02  Score=18.75  Aligned_cols=19  Identities=11%  Similarity=0.090
Sum_probs=0.0  Template_Neff=7.500

Q ss_pred              CcchhHHHHHHHHHhcccc
Q Q_5657867_1       1 MRHSAIGMLAVLVACREHP   19 (377)
Q Consensus         1 m~~~~~~~~~~~~~~~~~~   19 (377)
                      ||++.+..++.++++...+
T Consensus         1 Mk~~~~~~~l~l~~~s~~a   19 (96)
T ECOD_000005607    1 MKKRIIAAALLATVASFST   19 (96)
T ss_dssp             CCCCCCCCCCCCCCCCCCC
T ss_pred             ChHHHHHHHHHHHHHHHHH
```

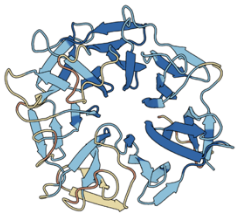

**Supplementary Figure 18:** Representative example of a novel undefined family (AFDB entry: A0A9D7CAU2). HHpred analysis yielded no significant propeller-fold hits. The predicted structure is colored according to pLDDT scores

```
Query         Q_A0A2N2ZE83_1
Match_columns 552
No_of_seqs    1238 out of 4157
Neff          12.7115
Searched_HMMs 30134
Date          Thu Sep 10 15:26:59 2026
Command       hhsearch -cpu 8 -i ../results/full.a3m -d /cluster/toolkit/production/databases/hh-suite/pfama/pfama -o ../
results/A0A2N2ZE83_1.hhr -oa3m ../results/A0A2N2ZE83_1.a3m -p 20 -Z 250 -loc -z 1 -b 1 -B 250 -ssm 0 -sc 1 -seq 1 -dbstrlen
10000 -norealign -maxres 32000 -contxt /cluster/toolkit/production/bioprogs/tools/hh-suite-build-new/data/context_data.crf

 No Hit                             Prob E-value P-value  Score    SS Cols Query HMM  Template HMM
  1 PF30506.1 ; Beta-prop_MIP2 ; M  99.5 1.2E-17 4.1E-22  153.0   0.0  234  113-377     4-306 (397)
  2 PF31099.1 ; SHKBP1_KCTD3_C ; S  99.4 1.8E-17 5.9E-22  153.1   0.0  346  110-524     3-384 (419)
  3 PF23381.2 ; Beta-prop_IFT122_1  99.4 2.5E-17 8.2E-22  142.1   0.0  258   72-372     1-273 (273)
  4 PF23798.2 ; Beta-prop_SPT8 ; S  99.4 5.6E-17 1.9E-21  149.7   0.0  310   69-422    11-420 (422)
  5 PF23383.2 ; Beta-prop_IFT140_1  99.4 6.5E-17 2.1E-21  148.4   0.0  300  111-451     9-406 (406)
  6 PF02239.23 ; Cytochrom_D1 ; Cy  99.4 6.5E-17 2.2E-21  147.9   0.0  324   78-441     1-394 (394)
  7 PF23586.3 ; Beta-prop_NWD2_C ;  99.4    9E-17    3E-21  144.6   0.0  302   64-414     2-355 (356)
  8 PF24017.3 ; Beta-prop_SCAP ; S  99.4 1.5E-16 4.9E-21  150.4   0.0  411   64-522     1-495 (495)
  9 PF25178.1 ; Beta-prop_WDR41 ;   99.3 2.2E-16 7.4E-21  146.5   0.0  363  122-533     2-439 (439)
 10 PF23798.2 ; Beta-prop_SPT8 ; S  99.3    5E-16 1.7E-20  143.2   0.0  340  115-497    11-420 (422)
 11 PF23586.3 ; Beta-prop_NWD2_C ;  99.3 5.8E-16 1.9E-20  139.3   0.0  287  194-523    10-354 (356)
 12 PF29630.1 ; ASA1 ; ASTRA-assoc  99.3 9.2E-16 3.1E-20  140.1   0.0  297   64-413     3-393 (393)
 13 PF24797.2 ; Beta-prop_WDR35_TU  99.3 1.1E-15 3.6E-20  136.1   0.0  266   69-374    13-333 (333)
 14 PF02239.23 ; Cytochrom_D1 ; Cy  99.3 1.3E-15 4.3E-20  139.2   0.0  308  163-515     1-394 (394)
 15 PF14655.13 ; RAB3GAP2_N ; Rab3  99.2 1.6E-15 5.4E-20  138.8   0.0  307   72-437     3-397 (397)
 16 PF21032.3 ; PROPPIN ; PROPPIN   99.2 1.8E-15    6E-20  135.4   0.0  309   72-424     1-339 (345)
 17 PF25391.2 ; WD40_Gbeta ; G pro  99.2 2.2E-15 7.3E-20  134.4   0.0  270   63-372    48-340 (340)
 18 PF24796.2 ; WDR55 ; WDR55       99.2 2.4E-15 7.9E-20  131.3   0.0  269   70-374     7-294 (296)
 19 PF23760.2 ; Beta-prop_DCAF12 ;  99.2 2.8E-15 9.2E-20  136.2   0.0  310   66-447     8-382 (383)
 20 PF26550.2 ; Tricorn_2nd ; Tric  99.2 3.4E-15 1.1E-19  132.6   0.0  296   73-416     1-333 (333)
```

**Supplementary Figure 19:** HHpred analysis result of the propeller domain A0A2N2ZE83 against the Pfam-A database.

```
Query         Q_A0A9D1M187_1
Match_columns 595
No_of_seqs    783 out of 2501
Neff          12.0898
Searched_HMMs 30134
Date          Thu Sep 10 15:22:39 2026
Command       hhsearch -cpu 8 -i ../results/full.a3m -d /cluster/toolkit/production/databases/hh-suite/pfama/pfama -o ../
results/A0A9D1M187_1.hhr -oa3m ../results/A0A9D1M187_1.a3m -p 20 -Z 250 -loc -z 1 -b 1 -B 250 -ssm 0 -sc 1 -seq 1
-dbstrlen 10000 -norealign -maxres 32000 -contxt /cluster/toolkit/production/bioprogs/tools/hh-suite-build-new/data/
context_data.crf

 No Hit                             Prob E-value P-value  Score    SS Cols Query HMM  Template HMM
  1 PF16465.11 ; DUF5046 ; Domain   97.1 1.5E-06 4.9E-11   72.2   0.0   49  511-561   204-255 (264)
  2 PF16465.11 ; DUF5046 ; Domain   96.0 0.00012   4E-09   61.0   0.0  188  229-425    30-254 (264)
  3 PF16465.11 ; DUF5046 ; Domain   95.9 0.00017 5.5E-09   60.2   0.0  143   52-200    97-254 (264)
  4 PF16465.11 ; DUF5046 ; Domain   95.4 0.00054 1.8E-08   57.2   0.0  209   47-284    16-252 (264)
  5 PF14903.12 ; WG_beta_rep ; WG   94.5  0.0025 8.2E-08   34.4   0.0   35  513-548     1-35  (35)
  6 PF14903.12 ; WG_beta_rep ; WG   90.9   0.034 1.1E-06   29.4   0.0   35  152-188     1-35  (35)
  7 PF14903.12 ; WG_beta_rep ; WG   89.7   0.056 1.9E-06   28.4   0.0   35  378-413     1-35  (35)
  8 PF29747.1 ; TP_0783 ; TP_0783   86.5    0.14 4.6E-06   46.7   0.0  173    9-198     1-188 (358)
  9 PF14903.12 ; WG_beta_rep ; WG   83.6    0.25 8.2E-06   25.6   0.0   29   65-93      1-29  (35)
 10 PF08139.18 ; LPAM_1 ; Prokaryo  83.1    0.27 8.9E-06   21.9   0.0   18   15-32      1-18  (18)
 11 PF29792.1 ; HI_1099 ; Uncharac  78.4    0.52 1.7E-05   33.6   0.0   20   13-32      1-20  (103)
 12 PF04202.18 ; Mfp-3 ; Foot prot  76.9    0.63 2.1E-05   30.2   0.0   25   15-39      1-25  (71)
 13 PF16383.11 ; DUF4992 ; Domain   76.8    0.63 2.1E-05   37.4   0.0   28    7-34      1-28  (183)
 14 PF10795.15 ; DUF2607 ; Protein  72.4    0.99 3.3E-05   31.8   0.0   31    4-34      2-32  (97)
 15 PF24052.3 ; Phage_lipoprotein   71.9       1 3.5E-05   29.9   0.0   23   12-34      1-23  (75)
 16 PF13956.12 ; Ibs_toxin ; Toxin  69.9     1.2 4.1E-05   19.4   0.0   17   14-30      1-17  (18)
 17 PF12930.13 ; DUF3836 ; Family   62.9     2.1 7.1E-05   32.1   0.0   27    9-35      1-27  (129)
 18 PF30535.1 ; Nlp-28-30 ; Neurop  61.9     2.3 7.6E-05   27.4   0.0   21   15-35      1-24  (70)
 19 PF30314.1 ; TP013X ; TP013X li  59.3     2.7 9.1E-05   39.3   0.0   31    4-34      1-31  (409)
 20 PF16390.12 ; DUF4999 ; Domain   58.5     2.9 9.6E-05   27.3   0.0   24   11-34      1-24  (75)
```

**Supplementary Figure 20:** HHpred analysis result of the propeller domain A0A9D1M187 against the Pfam-A database.

```
Query         Q_A0A9D7N7Y6_1
Match_columns 622
No_of_seqs    543 out of 4591
Neff          10.6643
Searched_HMMs 30134
Date          Thu Sep 10 15:23:31 2026
Command       hhsearch -cpu 8 -i ../results/full.a3m -d /cluster/toolkit/production/databases/hh-suite/pfama/pfama
-o ../results/A0A9D7N7Y6_1.hhr -oa3m ../results/A0A9D7N7Y6_1.a3m -p 20 -Z 250 -loc -z 1 -b 1 -B 250 -ssm 0 -sc 1 -seq
1 -dbstrlen 10000 -norealign -maxres 32000 -contxt /cluster/toolkit/production/bioprogs/tools/hh-suite-build-new/
data/context_data.crf

 No Hit                             Prob E-value P-value  Score    SS Cols Query HMM  Template HMM
  1 PF27095.1 ; Conj_beta_prop_9 ;  98.7 5.5E-12 1.8E-16  128.8   0.0  184  311-502   209-415 (417)
  2 PF27095.1 ; Conj_beta_prop_9 ;  98.6 1.6E-11 5.3E-16  125.3   0.0  142   75-218    50-226 (417)
  3 PF01011.29 ; PQQ ; Glucose/eth  98.3 6.7E-10 2.2E-14  118.2   0.0  129  126-254    15-201 (583)
  4 PF13360.14 ; PQQ_2 ; Outer mem  98.0 7.4E-09 2.4E-13   97.2   0.0  195  275-473     2-259 (259)
  5 PF13360.14 ; PQQ_2 ; Outer mem  98.0 7.4E-09 2.5E-13   97.2   0.0  202   49-253     1-259 (259)
  6 PF01011.29 ; PQQ ; Glucose/eth  98.0 9.3E-09 3.1E-13  109.3   0.0  317   47-364    13-563 (583)
  7 PF13360.14 ; PQQ_2 ; Outer mem  97.9 1.1E-08 3.7E-13   96.0   0.0  197  127-325     2-259 (259)
  8 PF27095.1 ; Conj_beta_prop_9 ;  97.9 1.1E-08 3.7E-13  104.0   0.0  269   41-319   104-417 (417)
  9 PF13360.14 ; PQQ_2 ; Outer mem  97.8 3.4E-08 1.1E-12   92.6   0.0  196  202-397     2-259 (259)
 10 PF27095.1 ; Conj_beta_prop_9 ;  97.7 5.1E-08 1.7E-12   99.1   0.0  242  263-510    49-343 (417)
 11 PF01011.29 ; PQQ ; Glucose/eth  97.4 3.6E-07 1.2E-11   97.0   0.0  281  229-510     7-497 (583)
 12 PF27482.1 ; Beta-prop_YycP ; Y  97.2 9.4E-07 3.1E-11   87.4   0.0  235  127-365    22-334 (351)
 13 PF27482.1 ; Beta-prop_YycP ; Y  96.9 4.5E-06 1.5E-10   82.5   0.0  236  238-477    22-334 (351)
 14 PF25293.2 ; Beta-prop_EMC1_N ;  96.9 5.3E-06 1.8E-10   82.2   0.0  302  170-474     3-382 (383)
 15 PF05567.18 ; T4P_PilY1 ; PilY1  96.7   1E-05 3.4E-10   79.8   0.0  169  331-501     7-302 (338)
 16 PF01011.29 ; PQQ ; Glucose/eth  96.7 1.3E-05 4.5E-10   84.7   0.0  209  301-510     7-330 (583)
 17 PF13570.14 ; Beta-prop_ACSF4 ;  96.6 1.5E-05 4.9E-10   77.7   0.0  271  193-465    17-351 (352)
 18 PF27482.1 ; Beta-prop_YycP ; Y  96.6 2.1E-05 6.8E-10   77.6   0.0  238   49-291    21-331 (351)
 19 PF25293.2 ; Beta-prop_EMC1_N ;  96.2 6.2E-05   2E-09   74.4   0.0  283   41-326    32-382 (383)
 20 PF05567.18 ; T4P_PilY1 ; PilY1  96.2 6.4E-05 2.1E-09   74.0   0.0  166  187-352     7-302 (338)
```

**Supplementary Figure 21:** HHpred analysis result of the propeller domain A0A9D7N7Y6 against the Pfam-A database.

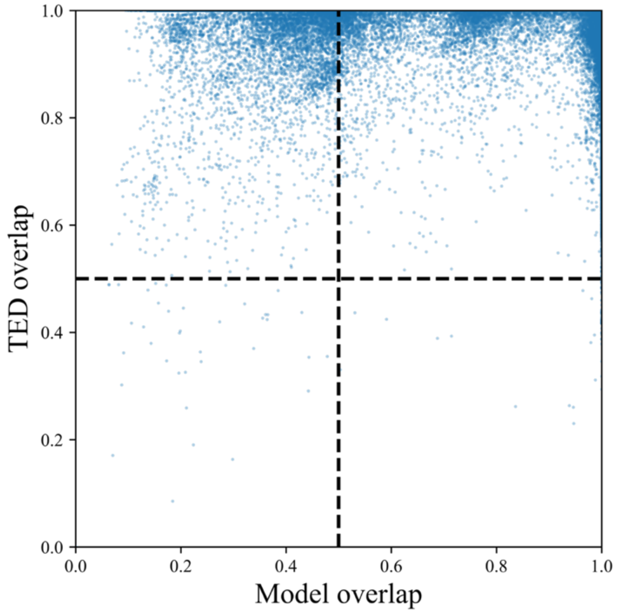


**Supplementary Figure 22:** Comparison of residue-level coverage between TREAD predictions and curated annotations for the same propeller proteins in AFDB. Propeller proteins detected by the TREAD model and independently curated in the TED database were compared at the residue level. For each protein, the fraction of residues covered by TED annotations that are also predicted by TREAD (TED overlap; y-axis) and the fraction of residues predicted by TREAD that are also annotated in TED (model overlap; x-axis) were computed. Each point represents one protein present in both datasets. The vertical (x = 0.5) and horizontal (y = 0.5) reference lines are shown for guidance.

```
Match_columns 238
No_of_seqs    158 out of 1480
Neff          9.56273
Searched_HMMs 143640
Date          Mon Dec  1 18:28:20 2025
Command       hhsearch -cpu 8 -i ../results/full.a3m -d /cluster/toolkit/production/databases/hh-suite/ECOD/ECOD_F70_20230309 -d /cluster/
toolkit/production/databases/hh-suite/mmcif70/pdb70 -o ../results/1268068_2.hhr -oa3m ../results/1268068_2.a3m -p 20 -Z 250 -loc -z 1 -b 1 -B
250 -ssm 2 -sc 1 -seq 1 -dbstrlen 10000 -norealign -maxres 32000 -contxt /cluster/toolkit/production/bioprogs/tools/hh-suite-build-new/data/
context_data.crf

 No Hit                             Prob E-value P-value  Score    SS Cols Query HMM  Template HMM
  1 7TPU_A Beta-lactamase domain-c  98.7 3.7E-05 2.6E-10   73.1  25.3  225    5-237     2-247 (616)
  2 7TPU_A Beta-lactamase domain-c  98.0  0.0041 2.9E-08   59.8  20.5  180   53-236     1-198 (616)
  3 7TPU_A Beta-lactamase domain-c  98.0   0.016 1.1E-07   55.8  24.3  199    1-204    50-266 (616)
  4 7TPU_A Beta-lactamase domain-c  96.9    0.11   8E-07   50.3  14.7  134   99-238     1-145 (616)
  5 ECOD_000118973_e1k28D4 | 3071.  21.3 3.6E+02  0.0025   19.0   2.5   27   43-69     22-48  (64)

No 1
>7TPU_A Beta-lactamase domain-containing protein; polyglycine hydrolase, chitinase-modifying protein, serine protease, beta lactamase-like,
HYDROLASE; HET: BMA, MAN, NAG; 2.194A {Fusarium vanettenii}
Probab=98.75  E-value=3.7e-05  Score=73.13  Aligned_cols=225  Identities=10%  Similarity=0.070  Sum_probs=141.4  Template_Neff=10.800

Q ss_pred             EEEEEeCCCCCceEEEEecCCcccccHHHHHHHHHcCceeEEEEEc----CCeEEEEEeCCCCCeeEEEEeCCCCc--HH
Q Q_1268068_2    5 LIVVTQQETGMYNQTWFYGGSGNSLQEDKIKEYWNEDFYINSVAYT----SKGWFVTMAKGLKWTNQSYSYKSSWP--DE   78 (238)
Q Consensus      5 w~~v~~~~~~~~~Q~~~~~~~~~~~~~~~I~~~w~~g~~It~i~~~----~~~w~~v~s~~~~~~~Q~~~~~~~~~~~--~~   78 (238)
                   |..++..+..-..-.+.+.....+| ...+++...+||++.+|...    ...+.+|+.+.++ ..+.+....+..  .+
T Consensus      2 ~~~~~~~~~~~~~~~~~~~~~~~~~-~~~~~~~~~~g~~~~~~~~~~~~~~~~~~v~~~~~g-~~~~~~~~~~~~~~~~~   79 (616)
T 7TPU_A         2 FLPNQRRSNVTSHVETYYSVDGATH-AEKSKALKADGYRIVSLSSYGSPDSANYAAIWVQEEG-PSFEIIHDADEATYNS   79 (616)
T ss_dssp          -----------CCEEEEEEEECHHHH-HHHHHHHHHTTEEEEEEEEESCGGGCEEEEEEEEECCC-CCEEEEEEEECHHHHHH
T ss_pred             CCCcccCCCCCCCeEEEecCCHHHH-HHHHHHHHhCCeEEEEEEecCCCCCceEEEEEEecCC-CCeEEEEeCCHHHHHH
Confidence       44555555443333444443233444 366777888999999999873    4568889988765 233333332222  45


Q ss_pred             HHHHHHHcCcEEEEEEEc----CCeEEEEEEeCCCCCcceeEe-C-ChhHHHHHHHHHHHcCceEEEEEE----eCCEEE
Q Q_1268068_2   79 WIQEKRKSGYMITSLSTS----GSNWMVVMSKNTDYKTQEICS-A-PWSTMKDWIKKWWNNDYYITSLTC----RNGMWT  148 (238)
Q Consensus     79 ~I~~~w~~g~~It~~~~~~----~~~w~vv~sk~~~~~~Q~~~~-~-~~~~~~~~~I~~~w~~G~~It~i~~----~~~~w~  148 (238)
                   .+.+..++||++.++...    ...|.+|+.+..... ..+.. . +...+.+.+.   ++|+++.++..    +...+.
T Consensus     80 ~~~~~~~~G~~~~~~~~~~~~~g~~~~~~i~~~~~~~~~~-~~~~~~~~~~~~~~~~~~~~~---~~g~~~~~~~~~~~~g~~~y~  155 (616)
T 7TPU_A        80 WLQTWKSRGYVSTQVSATGPAENAVFAGVMENINVAN-WFQSCELENPWAFSNTTG---NVDVVVKGFRMFGTPEERRYC  155 (616)
T ss_dssp          HHHHHHTTTCEEEEEEEEESCGGGCEEEEEEEECCCCSC-EEEEEEESSGGGHHHHHT---TCCEEEEEEEEEECSSSEEEE
T ss_pred             HHHHHHhCCceeEEEEEEecCCCcEEEEEEEeCCCCc-cEEEeCCCCHHHHHhhhc---CCCceEEEEEEeCCCCCcEEE
Confidence       5667778899999999864    347999999855432 22222 3 4455555554   89999999984    234788


Q ss_pred             EEEECCCCCeEEEEEEEec--CchhhHHHHHHHHHHcC-cEEEEEEEeC-CEEEEEEEccCCCCCcc-eEEEEcccChHHHH
Q Q_1268068_2  149 VVMSKTSLYIDQSYMSSS--TTSGIKEKIKKKWEEG-YRIIAFEFGG-GEYLCVMCKLAGNKTPM-QSYQIEPSDVSGFI  223 (238)
Q Consensus    149 vV~s~~~~~~~~~Q~~~~~~~~--~~~~~~~~~~~i~~~w~~g-y~It~i~~~~~-~~w~vv~sk~~~~~~~~~-q~~~~~~~~~~~~~i  223 (238)
                   +|+-.+.+...+.+.+..  +..++...+.+...+| +++..+...+ .++..++.+...  ... ..+..+........
T Consensus    156 ~vw~~~~~~~~~~~~~~~~~~~~~~~s~~~~~~~~~~~~~~~~~g~~r~~~~~~~~g~~~~~~v~~~~~~--~~~~~~~~~~~~~~~~~~~~  233 (616)
T 7TPU_A       156 ILGHENVGNEQTTIQYSTPSFTVNFASTFEAETTKRFWRPSRLFLSEDHIITPSFADTSV--GKWSHAVDLTKAELKEKI  233 (616)
T ss_dssp          EEEEECTTCCCEEECCCBTTBCCCHHHHHHHHHHSTTCEEEEEEEECTTCCEEEEEEECCCC--SSEEEEEEECHHHHHHH
T ss_pred             EEEEecCCCCceEEEecCCCccccHHHHHHHHHhcCCeeeEEEEEcCCCeEEEeEeeCCC--CceEEEecCCHHHHHHHH
Confidence       888887655556665555  5555688899999999 9999998664 468888887654  211 12222334567777
```

**Supplementary Figure 23:** HHpred analysis result of the five-copy barrel-like domain A0AAW5V244 against the ECOD70 and PDB70 database.

```
Match_columns 254
No_of_seqs    1183 out of 1185
Neff          13.1961
Searched_HMMs 71734
Date          Tue Dec  2 15:26:29 2025
Command       hhsearch -cpu 8 -i ../results/full.a3m -d /cluster/toolkit/production/databases/hh-suite/ECOD/ECOD_F70_20230309
-o ../results/2164156.hhr -oa3m ../results/2164156.a3m -p 20 -Z 250 -loc -z 1 -b 1 -B 250 -ssm 2 -sc 1 -seq 1 -dbstrlen 10000
-norealign -maxres 32000 -contxt /cluster/toolkit/production/bioprogs/tools/hh-suite-build-new/data/context_data.crf

 No Hit                             Prob E-value P-value  Score    SS Cols Query HMM  Template HMM
  1 ECOD_000401794_e3ow8B1 | 5.1.4  99.9 2.6E-23 3.7E-28  145.0  19.2  214    3-226    81-300 (300)
  2 ECOD_002801194_e7oopY1 | 5.1.5  99.9 2.6E-23 3.7E-28  145.0  19.2  214    3-226    81-300 (300)
  3 ECOD_001291899_e4ci8A2 | 5.1.4  99.9 4.9E-23 6.8E-28  145.7  20.2  219    6-231    36-285 (326)
  4 ECOD_001792730_e3jcrL2 | 5.1.4  99.9   2E-22 2.8E-27  140.8  22.8  214   10-233     2-229 (304)
  5 ECOD_001873820_e4j8bA1 | 5.1.4  99.9 1.3E-22 1.8E-27  141.3  21.6  243    2-253    31-281 (298)
  6 ECOD_002125194_e5oqlX1 | 5.1.4  99.9 1.4E-22 1.9E-27  143.6  21.0  239    2-253    37-292 (331)
  7 ECOD_002723796_e7luvE1 | 5.1.4  99.9 2.7E-22 3.7E-27  138.6  19.6  227    2-233    21-270 (282)
  8 ECOD_001792730_e3jcrL2 | 5.1.4  99.9 1.3E-22 1.9E-27  141.7  17.9  212    2-223    86-304 (304)
  9 ECOD_002817982_e7mqaLQ3 | 5.1.  99.9 1.6E-22 2.2E-27  141.1  18.1  221    2-233    65-303 (305)
 10 ECOD_002442391_e6f3tA1 | 5.1.5  99.9 3.4E-22 4.8E-27  143.1  20.1  222    2-233   105-332 (360)
 11 ECOD_002817977_e7mqaLO1 | 5.1.  99.9 7.7E-22 1.1E-26  139.0  21.5  241    3-253    33-298 (327)
 12 ECOD_002817936_e7mq8LO3 | 5.1.  99.9 9.5E-22 1.3E-26  139.0  21.6  241    3-253    33-310 (339)
 13 ECOD_002501370_e6nd4W1 | 5.1.5  99.9 3.2E-22 4.5E-27  144.9  19.4  219    2-233    25-248 (385)
 14 ECOD_002861541_e7somaa1 | 5.1.  99.9 8.7E-22 1.2E-26  136.6  20.7  239    2-253    32-276 (292)
 15 ECOD_002756041_e7mt1A1 | 5.1.4  99.9 8.6E-22 1.2E-26  138.3  20.9  239    2-253    39-303 (321)
 16 ECOD_002573113_e6tb4G2 | 5.1.5  99.9 1.3E-21 1.8E-26  140.6  21.6  222    2-233    93-342 (366)
 17 ECOD_002810373_e7ktrB1 | 5.1.5  99.9 7.9E-22 1.1E-26  139.6  20.2  218    2-229    34-289 (331)
 18 ECOD_000140824_e3ow8A1 | 5.1.4  99.9 5.4E-22 7.5E-27  137.8  18.9  215    2-226    73-293 (293)
 19 ECOD_002055685_e5mwjA1 | 5.1.4  99.9 3.8E-22 5.4E-27  141.6  18.4  214    2-226   108-327 (328)
 20 ECOD_002859845_e7s22A1 | 5.1.4  99.9 4.7E-22 6.5E-27  139.1  18.3  222    2-229    74-306 (308)
 21 ECOD_000002206_e1tbgA1 | 5.1.4  99.9 1.8E-21 2.5E-26  138.1  21.4  244    3-253    34-282 (340)
 22 ECOD_002323838_e5i2tA2 | 5.1.4  99.9   2E-21 2.7E-26  137.7  21.5  241    2-253    30-311 (340)
 23 ECOD_001756814_e3jb9K1 | 5.1.5  99.9 7.5E-21   1E-25  133.7  24.2  227   15-254     1-232 (317)
 24 ECOD_002069376_e5mzhA1 | 5.1.5  99.9 8.4E-22 1.2E-26  139.6  19.5  216    2-227   119-340 (340)
 25 ECOD_002321494_e3mzkA1 | 5.1.4  99.9 3.7E-22 5.1E-27  138.9  17.2  232    2-239    29-297 (297)
 26 ECOD_002859845_e7s22A1 | 5.1.4  99.9 1.1E-20 1.5E-25  132.3  24.6  228   15-253     3-244 (308)
 27 ECOD_001298457_e4owrA1 | 5.1.4  99.9 1.7E-21 2.4E-26  137.6  20.7  210   16-235     1-324 (335)
 28 ECOD_002832128_e7s23B1 | 5.1.4  99.9 6.6E-22 9.2E-27  138.5  18.4  221    2-228    74-311 (313)
 29 ECOD_000043699_e1vyhD1 | 5.1.4  99.9   4E-22 5.5E-27  139.8  17.1  218    2-229    37-280 (317)
 30 ECOD_002643920_e7bz2B1 | 5.1.4  99.9 2.4E-21 3.4E-26  137.0  21.2  244    3-253    28-276 (334)
 31 ECOD_002832128_e7s23B1 | 5.1.4  99.9   1E-20 1.4E-25  132.5  23.9  228   15-253     3-250 (313)
 32 ECOD_002388925_e6cvzA1 | 5.1.5  99.9 2.3E-21 3.2E-26  136.3  20.6  223    1-233    21-286 (321)
 33 ECOD_002142227_e6exnJ1 | 5.1.4  99.9   3E-21 4.2E-26  137.2  21.2  209   15-233    21-235 (342)
 34 ECOD_001884253_e5lj5J1 | 5.1.4  99.9 6.6E-21 9.2E-26  134.5  22.6  209   15-233    20-234 (327)
 35 ECOD_000002212_e1nr0A1 | 5.1.4  99.9 3.1E-21 4.3E-26  135.0  20.8  223    2-233    36-275 (311)
 36 ECOD_001495248_e4d5lg1 | 5.1.4  99.9 3.9E-21 5.4E-26  134.9  21.2  232   11-253     1-244 (314)
 37 ECOD_000002217_e1vyhC1 | 5.1.4  99.9 1.4E-21 1.9E-26  136.6  18.8  222    2-233    37-277 (310)
 38 ECOD_001872125_e5gmkO1 | 5.1.4  99.9 5.3E-21 7.4E-26  135.6  21.9  210   14-233    19-234 (337)
 39 ECOD_000146658_e3jroA1 | 5.1.4  99.9 7.2E-22   1E-26  136.8  17.1  231    2-238    22-289 (289)
 40 ECOD_002729429_e7edxE1 | 5.1.5  99.9 2.4E-21 3.3E-26  139.9  20.2  234    2-245   105-344 (382)
 41 ECOD_002905442_e7mq8LR1 | 5.1.  99.9 4.2E-22 5.9E-27  140.2  16.0  223    2-234    80-327 (327)
 42 ECOD_001556074_e4xfvA1 | 5.1.4  99.9 1.1E-21 1.5E-26  141.0  18.3  228    2-235    32-377 (378)
 43 ECOD_002066817_e5mzhB2 | 5.1.5  99.9 5.4E-21 7.5E-26  135.5  21.6  239    2-253    77-321 (342)
 44 ECOD_002772346_e7mdzgg1 | 5.1.  99.9 5.6E-21 7.8E-26  134.0  21.4  232   11-253     1-244 (313)
 45 ECOD_002321462_e4jxmA1 | 5.1.4  99.9 7.5E-21   1E-25  134.5  22.2  236    8-253     1-304 (336)
```

**Supplementary Figure 24:** HHpred analysis result of the decorated propeller-like domain A0A838VL09 against the ECOD70 database.

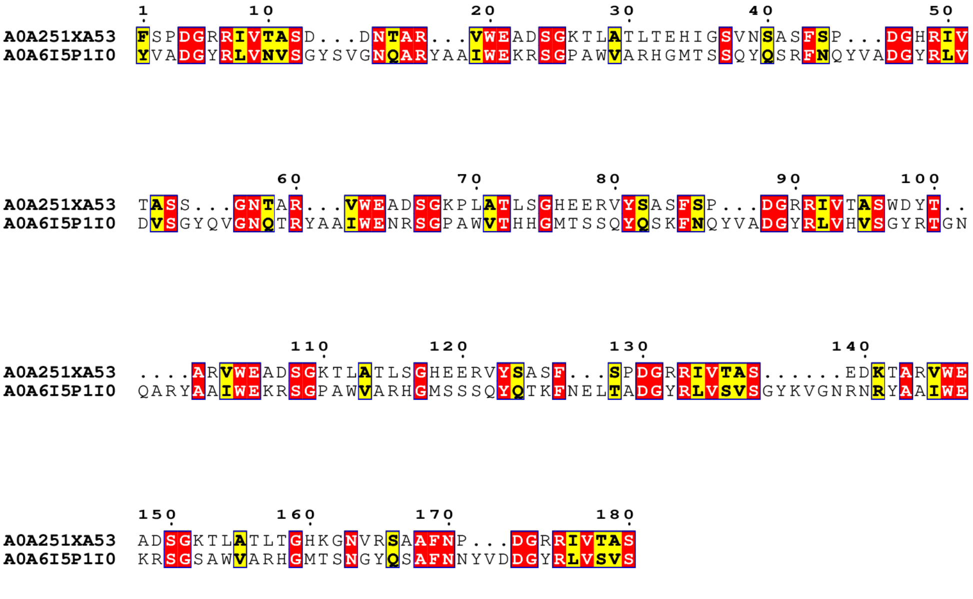


**Supplementary Figure 25:** Full-length pairwise alignment between a 12-bladed propeller domain (A0A251XA53) and a five-copy barrel-like domain (A0A6I5P1I0), as reported by BLASTp (E-value = 1e−13, 30% sequence identity, and 39% similarity). The alignment is visualized using ESPript 3.2, with identical and similar residues shaded in red and yellow, respectively.

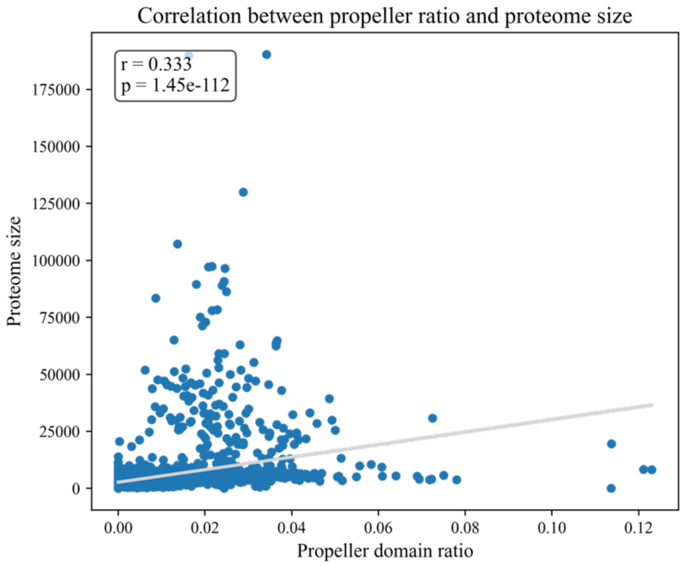


**Supplementary Figure 26:** Correlation between proteome size and the proportion of propeller-containing proteins across representative species included in the scan (Spearman R = 0.33, $p < 0.001$).

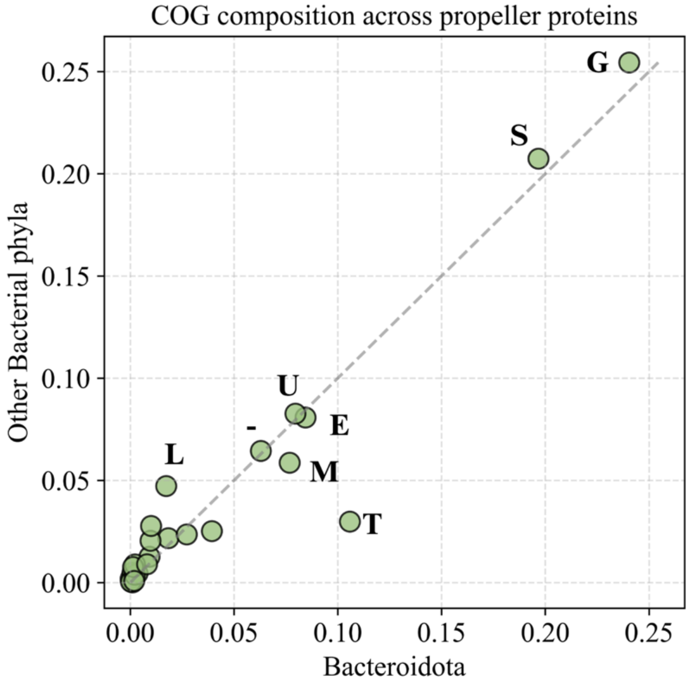


**Supplementary Figure 27:** Comparison of COG functional composition between propeller-containing proteins detected in Bacteroidota and those from other bacterial phyla. COG categories displaying pronounced compositional differences are labeled with the COG letter. G: Carbohydrate transport and metabolism; T: Signal transduction mechanisms; S/-: Function unknown; E: Amino acid transport and metabolism; U: Intracellular trafficking, secretion, and vesicular transport; M: Cell wall/membrane/envelope biogenesis; O: Posttranslational modification, protein turnover, chaperones; L: Replication, recombination and repair.

### Supplementary Tables

**Supplementary Table 1:**

| out_channel | hidden_dim | num_block | dropout | kernel_conv1 | kernel_block | f1 |
|---|---|---|---|---|---|---|
| 64 | 64 | 2 | 0.2 | 3 | 7 | 0.987787 |
| 64 | 64 | 5 | 0.2 | 3 | 3 | 0.987180 |
| 64 | 64 | 6 | 0.1 | 3 | 3 | 0.986746 |
| 64 | 64 | 3 | 0.2 | 11 | 7 | 0.986704 |
| 32 | 64 | 5 | 0.1 | 7 | 3 | 0.986563 |
| 32 | 64 | 3 | 0.1 | 11 | 7 | 0.986545 |
| 32 | 32 | 5 | 0.1 | 7 | 3 | 0.986325 |
| 64 | 32 | 5 | 0.1 | 3 | 7 | 0.986319 |
| 32 | 32 | 3 | 0.1 | 3 | 7 | 0.986286 |
| 32 | 64 | 2 | 0.1 | 11 | 11 | 0.986174 |

Top 10 hyperparameter configurations identified via grid search for the TREAD RepeatsDB model. The parameters evaluated include *out_channel*, representing the number of channels per residue throughout the network; *num_block*, indicating the total count of ResNet blocks; and *hidden_dim*, specifying the dimensionality of the penultimate fully connected layer. Convolutional kernel sizes are reported as *kernel_conv1* for the initial layer and *kernel_block* for the subsequent ResNet blocks. Finally, *Dropout* denotes the regularization probability applied across the architecture to optimize generalization.

**Supplementary Table 2:**

| Model | Representation | Residue-level F1-score |
|---|---|---|
| TREAD | ProtT5 embedding | 0.910±0.010 |
| Linear | ProtT5 embedding | 0.851±0.087 |
| TREAD | One-hot encoding | 0.752±0.174 |

Ablation study in the training of TREAD RepeatsDB model.

**Supplementary Table 3:**

| kernel_conv1 | kernel_block | num_block | out_channel | hidden_dim | F1 |
|---|---|---|---|---|---|
| 11 | 7 | 1 | 128 | 64 | 0.945808 |
| 11 | 3 | 1 | 64 | 64 | 0.945040 |
| 11 | 3 | 2 | 64 | 64 | 0.945019 |
| 11 | 3 | 1 | 64 | 32 | 0.944812 |
| 11 | 3 | 2 | 64 | 32 | 0.944811 |
| 7 | 7 | 1 | 64 | 64 | 0.944801 |
| 7 | 3 | 2 | 64 | 32 | 0.944726 |
| 7 | 7 | 2 | 32 | 64 | 0.944724 |
| 7 | 11 | 3 | 64 | 64 | 0.944660 |
| 7 | 3 | 1 | 32 | 64 | 0.944588 |

Top 10 hyperparameter configurations identified via grid search for the propeller blade model. The parameters evaluated include *out_channel*, representing the number of channels per residue throughout the network; *num_block*, indicating the total count of ResNet blocks; and *Hidden_dim*, specifying the dimensionality of the penultimate fully connected layer. Convolutional kernel sizes are reported as *kernel_conv1* for the initial layer and *kernel_block* for the subsequent ResNet blocks.

**Supplementary Table 4:**

| COG Code | Description |
| --- | --- |
| J | Translation, ribosomal structure and biogenesis |
| A | RNA processing and modification |
| K | Transcription |
| L | Replication, recombination and repair |
| B | Chromatin structure and dynamics |
| D | Cell cycle control, cell division, chromosome partitioning |
| Y | Nuclear structure |
| V | Defense mechanisms |
| T | Signal transduction mechanisms |
| M | Cell wall/membrane/envelope biogenesis |
| N | Cell motility |
| Z | Cytoskeleton |
| W | Extracellular structures |
| U | Intracellular trafficking, secretion, and vesicular transport |
| O | Posttranslational modification, protein turnover, chaperones |
| C | Energy production and conversion |
| G | Carbohydrate transport and metabolism |
| E | Amino acid transport and metabolism |
| F | Nucleotide transport and metabolism |
| H | Coenzyme transport and metabolism |
| I | Lipid transport and metabolism |
| P | Inorganic ion transport and metabolism |
| Q | Secondary metabolites biosynthesis, transport and catabolism |
| R | General function prediction only |
| S | Function unknown |

Description of COG codes.

**Supplementary Table 5:**

| COG Category | Eukaryote | Prokaryote | Bacteroidota |
| --- | --- | --- | --- |
| J | 3.3% | 0.3% | 0.2% |
| A | 3.4% | 0.2% | 0.1% |
| K | 3.9% | 2.1% | 1.8% |
| L | 1.8% | 4.1% | 1.7% |
| B | 2.3% | 0.1% | 0.1% |
| D | 4.7% | 1.2% | 0.9% |
| Y | 0.6% | 0.0% | 0.0% |
| V | 0.1% | 0.6% | 0.1% |
| T | 9.8% | 4.7% | 10.6% |
| M | 0.4% | 6.2% | 7.7% |
| N | 0.1% | 1.7% | 1.0% |
| Z | 4.3% | 0.9% | 0.8% |
| W | 1.4% | 0.1% | 0.2% |

| | | | |
|---|---|---|---|
| U | 6.2% | 7.9% | 8.0% |
| O | 5.8% | 2.6% | 2.7% |
| C | 0.3% | 2.9% | 3.9% |
| G | 2.5% | 24.5% | 24.0% |
| E | 0.6% | 8.1% | 8.4% |
| F | 0.1% | 0.4% | 0.1% |
| H | 0.1% | 0.1% | 0.1% |
| I | 0.9% | 0.4% | 0.3% |
| P | 0.7% | 0.9% | 0.2% |
| Q | 1.3% | 2.4% | 1.0% |
| R/S/- | 40.4+5.1% | 20.9+6.8% | 19.7+6.2% |

Functional enrichment analysis of identified propeller proteins, categorized by COG classifications across Eukaryotes, Prokaryotes, and Bacteroidota.
